# Fibronectin-dependent tissue mechanics regulate the translation of segmentation clock oscillations into periodic somite formation

**DOI:** 10.1101/808121

**Authors:** Patrícia Gomes de Almeida, Pedro Rifes, Ana Patrícia Martins-Jesus, Gonçalo G. Pinheiro, Raquel P. Andrade, Sólveig Thorsteinsdóttir

**Author notes:** Correspondence to: Sólveig Thorsteinsdóttir. The Novo Nordisk Foundation Center for Stem Cell Biology (DanStem), University of Copenhagen, 2200 Copenhagen N, Denmark (Pedro Rifes). These authors contributed equally (Raquel P. Andrade, Sólveig Thorsteinsdóttir).

## Abstract

Somitogenesis starts with cyclic waves of expression of segmentation clock genes in the presomitic mesoderm (PSM) and culminates with periodic budding of somites in its anterior-most region. How cyclic clock gene expression is translated into timely morphological somite formation has remained unclear. A posterior to anterior gradient of increasing PSM tissue cohesion correlates with increasing fibronectin matrix complexity around the PSM, suggesting that fibronectin-dependent tissue mechanics may be involved in this transition. Here we address whether the mechanical properties of the PSM tissue play a role in regulating the pathway leading to cleft formation in the anterior PSM. We first interfered with cytoskeletal contractility in the chick PSM by disrupting actomyosin-mediated contractility directly or via Rho-associated protein kinase function. Then we perturbed fibronectin matrix accumulation around the PSM tissue by blocking integrin-fibronectin binding or fibronectin matrix assembly. All four treatments perturbed *hairy1* and *meso1* expression dynamics and resulted in defective somitic clefts. A model is presented where a gradient of fibronectin-dependent tissue mechanics participates in the PSM wavefront of maturation by ensuring the correct spatio-temporal conversion of cyclic segmentation clock gene expression into periodic somite formation.

## Introduction

Cells in the developing embryo are constantly receiving and integrating information, including mechanical signals generated by the adhesion to neighbor cells and/or the surrounding extracellular matrix (ECM). Cell-cell adhesion molecules and cell-ECM receptors, such as cadherins and integrins, respectively, are linked to the intracellular actomyosin cytoskeleton via intermediate proteins (Campbell and Humphries, 2011; Charras and Yap, 2018; Takeichi, 2014; Wolfenson et al., 2013). These adhesion complexes, called adhesomes, allow cells to perceive and respond to changes in their physical surroundings (Horton et al., 2016; Zaidel-Bar, 2013). Signaling events in adhesomes can impact the actomyosin cytoskeleton through the phosphorylation of non-muscle myosin II (NM II) which binds to actin and converts ATP into mechanical energy (Zaidel-Bar et al., 2015). The resulting actomyosin contractility leads to changes in cell shape and can transmit signals from integrin adhesomes to cadherin adhesomes and vice versa, as well as from the cell surface to the nucleus (Burute and Thery, 2012; Mui et al., 2016; Wolfenson et al., 2019). In this way, through continuous probing of their mechanical environment, cells adjust their shape, functions and behaviors, such as proliferation, differentiation, cell polarity and migration (Burute and Thery, 2012; Mui et al., 2016; Wolfenson et al., 2019). While morphogens have been extensively studied as major chemical regulators of developmental processes (Marek and Kubícek, 1981; Slack, 1987; Tiedemann, 1976), the importance of mechanical forces in embryo development has, until recently, received less attention (Marek and Kubícek, 1981; Slack, 1987; Tiedemann, 1976). It is, however, becoming increasingly clear that the ability of cells to sense and respond to mechanical signals regulates numerous basic developmental processes (e.g. Barriga et al., 2018; Brunet et al., 2013; Hiramatsu et al., 2013; Smutny et al., 2017).

One of the most conspicuous morphogenetic events during early vertebrate embryogenesis is the formation of somites, which are the source of axial skeleton and skeletal muscle precursor cells (Christ et al., 2007). Somites are spheres of epithelioid cells that are formed periodically from the anterior portion of the mesenchymal presomitic mesoderm (PSM), bilateral to the axial structures (Bailey and Dale, 2015). Temporal control of somite formation is dependent on cyclic waves of expression of segmentation clock genes, many of which are targets of the Notch signaling pathway (Dequéant et al., 2006; Masamizu et al., 2006; Palmeirim et al., 1997). These waves periodically sweep the PSM in a posterior to anterior direction (Aulehla and Pourquié, 2010; Bailey and Dale, 2015) and, as they reach the anterior PSM, oscillations slow down and then arrest (Morimoto et al., 2005; Shih et al., 2015). The transcription factor Mesp2/Meso1 is upregulated downstream of the segmentation clock in the anterior PSM, leading to Eph/Ephrin signaling and somitic cleft formation (Barrios et al., 2003; Nakajima et al., 2006; Saga, 2012; Watanabe et al., 2009), followed by progressive cell rearrangements into a somite (Martins et al., 2009; Morimoto et al., 2005; Shih et al., 2015).

Fibronectin is essential for somite formation in all vertebrate models studied to date (George et al., 1993; Georges-Labouesse et al., 1996; Goh et al., 1997; Koshida et al., 2005; Kragtorp and Miller, 2007; Rifes et al., 2007; Sato et al., 2007). Fibronectin matrix assembly is a complex cell-dependent process that requires the engagement and unfolding of globular fibronectin by the major fibronectin matrix assembly receptor, the α5β1 integrin, followed by fibrillogenesis involving fibronectin-fibronectin binding (Mao and Schwarzbauer, 2005; Singh et al., 2010). In the chick, a fibronectin matrix starts being assembled around the caudal PSM tissue and then gets progressively denser as the tissue matures (Rifes et al., 2007; Rifes and Thorsteinsdóttir, 2012). This results in the formation of a gradient of fibronectin matrix complexity along the PSM (Rifes and Thorsteinsdóttir, 2012), which correlates with a posterior to anterior gradient in cell density (Bénazéraf et al., 2010; Lawton et al., 2013; Mongera et al., 2018). At the rostral end, fibronectin is required for the polarization of N-cadherin and epithelialization of peripheral cells to form a somite (Martins et al., 2009; Rifes et al., 2007). Interestingly, adhesion to a fibronectin substrate was noted as a regulator of the oscillations of the segmentation clock gene *Lfng* in cultured mouse tailbud cells (Hubaud et al., 2017; Lauschke et al., 2013). Cell adhesion to fibronectin was linked to dampening and eventual arrest of *Lnfg* oscillations (Hubaud et al., 2017), reminiscent of what is observed in the anterior PSM prior to somite epithelialization. However, whether the mechanical properties of the PSM tissue play a role in the slowing down of segmentation clock oscillations and their conversion into segments remains unknown.

In this study, we addressed the involvement of PSM tissue mechanics in the regulation of both segmentation clock gene expression dynamics and subsequent somite formation using the chick embryo as a model. First, we experimentally perturbed actomyosin contractility by blocking either NM II ATPase activity or Rho-associated protein kinase (ROCK). We then addressed the role of the fibronectin matrix surrounding the PSM by blocking integrin-fibronectin binding through RGD or by perturbing extracellular fibronectin fibrillogenesis. We found that each one of the four treatments resulted in abnormal segmentation clock oscillations, mis-positioning of *meso1* expression in the rostral PSM and perturbations in somite morphogenesis. These results strongly suggest that fibronectin-dependent PSM tissue mechanics play a role in converting segmentation clock oscillations into periodic somite formation.

## Results

### Intracellular actomyosin contractility is required for timely segmentation clock oscillations and *meso1* activation

In the chick embryo, sequential pairs of somites bud off from the anterior PSM every 90 min, which corresponds to the period of segmentation clock oscillations (Figure 1 A). To investigate the involvement of intracellular actomyosin contractility in this process, the expression of the segmentation clock gene *hairy1* (Palmeirim et al., 1997) was analyzed in the PSM of embryo half explants cultured in the presence of either Blebbistatin, which directly inhibits the ATPase activity of NM II and consequently all actomyosin contractility (direct inhibition), or RockOut, a chemical inhibitor of ROCK I and II (ROCK I/II) enzymes involved in activating NM II (indirect inhibition) (Figure 1 B, C; Ringer et al., 2017; Straight et al., 2003; Yarrow et al., 2005). The contralateral control sides were cultured with an equal volume of DMSO.

**Figure 1.**
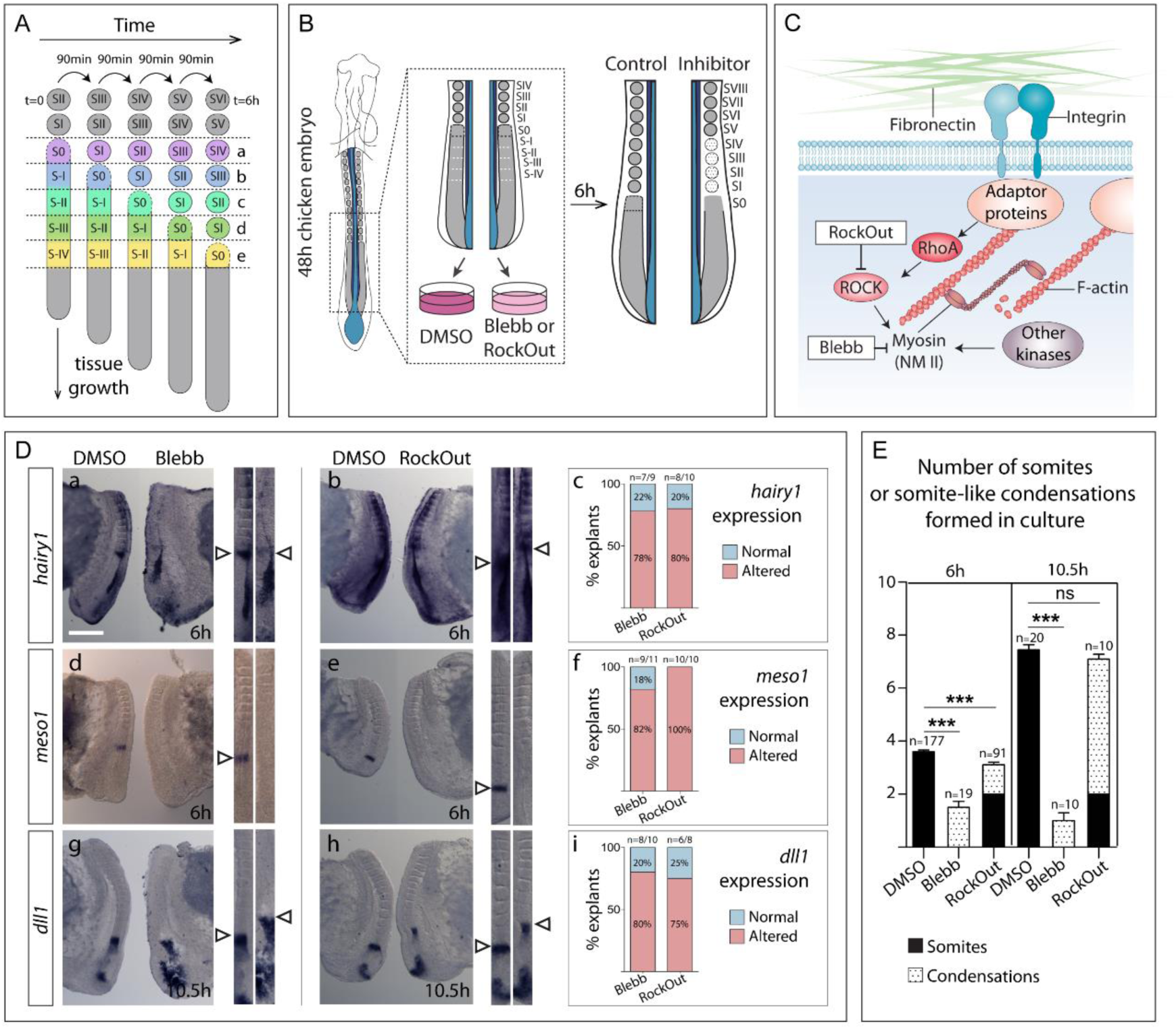
Blocking NM II or ROCK I/II activity leads to misregulation of *hairy1*, *meso1* and *dll1* expression and to defects in somite formation. (**A**) Schematic representation of chick PSM maturation and somite formation over time. A new pair of somites buds off from the anterior PSM every 90 min. This is also the period of segmentation clock oscillations in the chick embryo. With each new pair of somites, the previously formed somites mature (SI becomes SII, SII becomes SIII, etc). (**B**) Schematic representation of the explant culture system. Posterior explants of HH11-14 chick embryos were bisected along the midline and cultured for 6 (or 10.5) hours. One side of the explant was cultured with either Blebbistatin (Blebb) or RockOut, while the contra-lateral half was cultured with an equal volume of DMSO. **(C)** Schematic representation of the action of Blebbistatin and RockOut. Blebbistatin inhibits NM II ATPase activity directly while RockOut inhibits ROCK I/II-mediated phosphorylation of myosin light chain, thus indirectly decreasing NM II ATPase activity. (**D**) *In situ* hybridization for *hairy1* (a, b) *meso1* (d, e) and *dll1* (**g, h**) after 6 (a, b, d, e) or 10.5 hours of culture (g, h) in Blebbistatin (Blebb; a, d, g) and in RockOut-containing (b, e, h) media. Straightened images of the respective explant pairs (right) were aligned by SIV. Rostral is on top. Percentage of Blebbistatin-or RockOut-treated explants with altered *hairy1*, *meso1* and *dll1* expression compared to the contralateral controls is shown in c, f and i, respectively. **(E)** Number of somites (black bars) or somite-like condensations (dotted bars) formed in cultured explants. Explants cultured with DMSO formed sharp somite boundaries and clearly individualized somites. Blebbistatin-treated explants only formed 1-2 somite-like condensations. In RockOut treated explants the first two somites were normal while the remaining ones were cell condensations with poorly defined boundaries. p values were calculated using a paired Student’s t-test. ns – not significant, *** – p<0.01. Scale bar in D: 500 µm. Bars - standard error of the mean.

Explants cultured for 6 hours in each experimental condition presented significantly altered *hairy1* expression. *hairy1* expression was either absent or in a different phase of the cycle relative to the contralateral control in 80% of the Blebbistatin-(n=7/9) and RockOut-treated explants (n= 8/10; Figure 1 D, a-c), suggesting that temporal control of *hairy1* oscillations requires the generation of tensional cues mediated by NM II and ROCK I/II activity.

Segmentation clock oscillations are required for the correct spatial and temporal upregulation of *Mesp2* in the anterior PSM (Niwa et al., 2011; Saga and Takeda, 2001; Sato et al., 2002), which regulates downstream targets needed for the formation of the somitic cleft (Saga, 2012). The expression of the chick *Mesp2* homolog, *meso1*, was altered in Blebbistatin-(n=9/11) and RockOut-treated explants (n=10/10; Figure 1 D, d-f). *meso1* expression was either absent, located more rostrally or presented a different number of bands of expression, clearly indicating that the normal cycles of activation and suppression of *meso1* in the rostral PSM were altered. Importantly, *meso1* expression was also perturbed after 3 hours in culture with Blebbistatin or RockOut (n=8/9 and 5/5, respectively; Supplementary Figure 1 A-B), corresponding to an effect within two segmentation clock cycles. Furthermore, timely downregulation of *dll1* in the anterior-most PSM (Palmeirim et al., 1998), which normally occurs downstream of Meso1/Mesp2 activity (Takahashi et al., 2000; Takahashi et al., 2003), was not observed in either Blebbistatin-(80%, n=8/10) or RockOut-treated (75%, n=6/8) explants after 10.5h of culture (Figure 1 D, g-i). Together these data indicate that interfering with actomyosin contractility perturbs three sequential events: the spatio-temporal expression dynamics of *hairy1*, timely *meso1* expression and the downregulation of *dll1* expression in the anterior-most PSM.

Alterations in somite formation were observed concomitantly with the perturbations in *hairy1* and *meso1* expression. Control explants formed an average of 3.6 somites after 6 hours, consistent with a 90 min periodicity (Figure 1 E), while contralateral RockOut-treated explants formed 3.1 somites, of which only the first two somites were clearly individualized, while subsequent somite-like condensations were poorly defined (Figure 1 E). After 10.5 hours, control explants had formed an average of 7.5 somites, while RockOut-treated explants formed 7.1 somites of which the first two appeared normal, but the remaining ones were ill-defined (Figure 1 E). Importantly, explants cultured with Blebbistatin were unable to form more than 1-2 somite-like aggregates after 6 hours, or even 10.5 hours, of culture (Figure 1 E), evidencing an absolute requirement for NM II ATPase activity in somite formation. These effects were not due to an increase in apoptosis (Supplementary Figure 2 A-D).

Our data reveal a previously unknown role for NM II-and ROCK I/II-mediated cell contractility in the temporal regulation of the segmentation clock, *meso1* expression, *dll1* downregulation and, consequently, in somite formation.

### NM II activity is required for somite cleft formation and cell polarization

Somite formation involves a mesenchymal-to-epithelial transition (MET) of anterior PSM cells (Martins et al., 2009; Saga, 2012). To determine to what extent this process is impaired upon NM II or ROCK I/II inhibition, we performed a detailed analysis of the morphology of S0 to SIII in explants after a 6 hour culture period (regions e-b in Figure 1 A).

In control explants, S0 showed apically enriched N-cadherin (Figure 2 A, D) and some *zonula occludens* protein 1 (ZO-1) accumulation was observed apically (Figure 2 B, D, arrowheads). Peripheral cell alignment occurred (Figure 2 C, D) and fibronectin matrix was detected in the nascent somitic clefts (Figure 2 E, F, arrows). In contrast, in explants cultured with Blebbistatin, N-cadherin was homogeneous (Figure 2 G, J), ZO-1 immunostaining was absent (Figure 2 H, J) and neither peripheral cell alignment (Figure 2 I, J) nor fibronectin matrix accumulation within the tissue was observed (Figure 2 K, L). Furthermore, the continuous and dense fibronectin matrix observed surrounding the rostral PSM in control explants was disrupted in Blebbistatin-treated explants (compare Figure 2 E and K, arrowheads). Moreover, the characteristic nuclear alignment and F-actin apical enrichment observed in control SI (Supplementary Figure 3 A-C) was absent in Blebbistatin-treated explants and no signs of somitic boundaries could be detected (Supplementary Figure 3 D-F). We conclude that exposure of the S-IV and S-III regions of the PSM (regions e and d in Figure 1 A) to Blebbistatin for 6 hours completely blocks their capacity to form somites.

**Figure 2.**
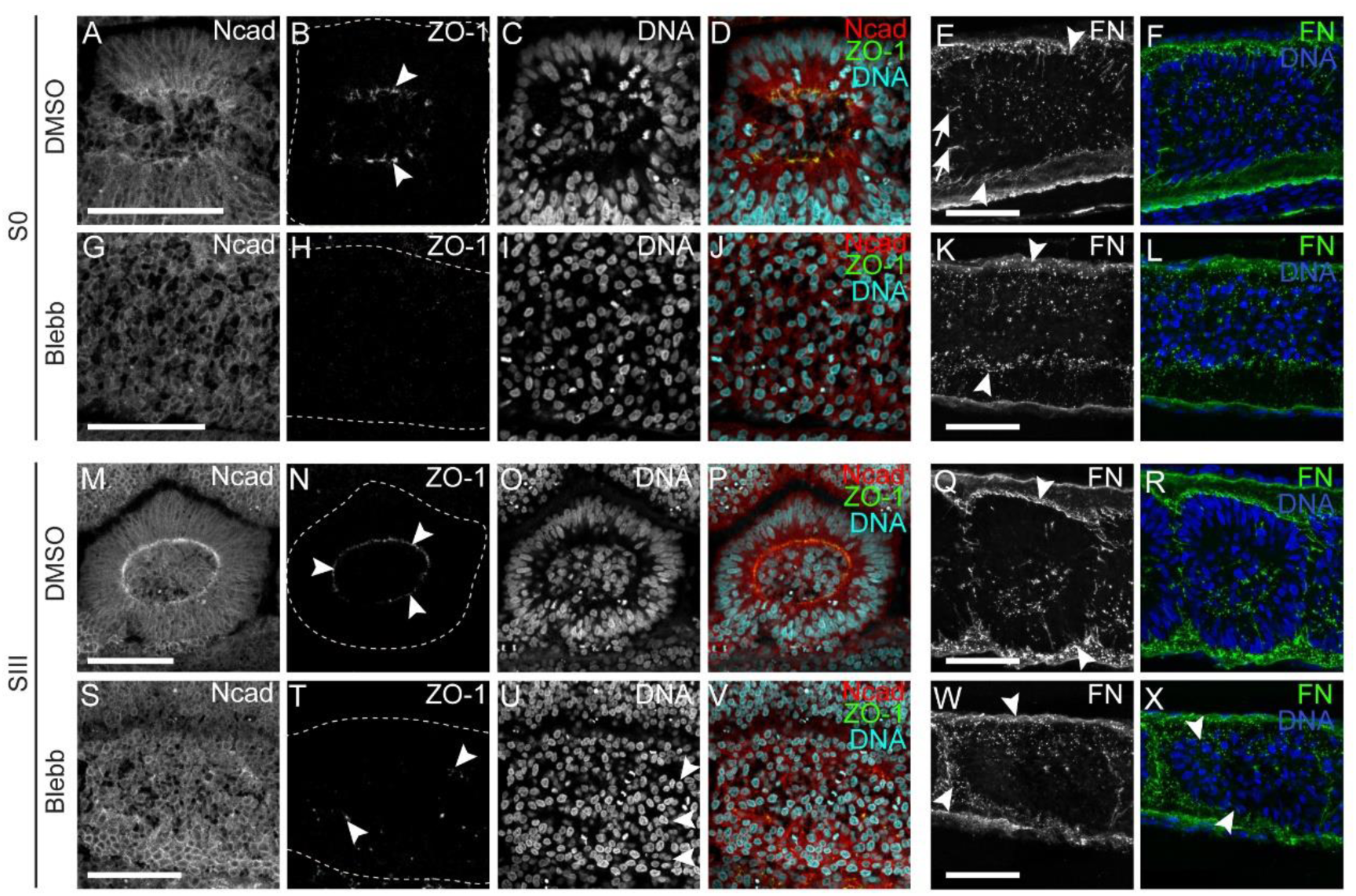
NM II inhibition abolishes N-cadherin and ZO-1 polarization and impairs fibronectin fibrillogenesis. (**A-X**) Sagittal optical sections of control explants (A-F, M-R) and their Blebbistatin-treated contralateral halves after 6 hours of culture, immunostained for N-cadherin (Ncad), ZO-1, fibronectin (FN), stained for DNA and imaged at S0 (A-L; region e in Figure 1 A) and SIII (M-X; region b in Figure 1 A) levels. S0 of DMSO-treated explants shows apically enriched N-cadherin (A) and ZO-1 (B, arrowheads) and peripheral nuclei are aligned (C, D), while no signs of polarized cell-cell adhesions (G, H) or nuclear alignment (I, J) are found in the contralateral Blebbistatin-treated explant. Apical polarization of N-cadherin and ZO-1 is maintained in SIII cells of DMSO-treated explants (M, N, arrowheads), but no polarized N-cadherin (S) or ZO-1 (T, arrowheads) are observed in contralateral Blebbistatin-treated halves. Nuclear alignment (U, arrowheads) and fibronectin assembly around somites (K, W, arrowheads) are also deficient in Blebbistatin-treated explants compared to contralateral controls (E, Q, arrowheads). Arrows in E show fibronectin assembly in the nascent somitic cleft. Rostral to the left and dorsal on top. Dashed lines mark borders of S0 (B, H) and SIII (N, T). FN – fibronectin. Ncad – N-cadherin. ZO-1 – *Zonula occludens* 1. Blebb – Blebbistatin. Scale bars: 50 µm.

We next turned our attention to SII and SIII somites after 6 hours of culture. These were at stage S-II and S-I in the PSM, respectively (regions c and b in Figure 1 A), when the explants were placed in culture and had thus already upregulated *meso1* (Buchberger et al., 1998). As before, in the presence of Blebbistatin, apical enrichment of N-cadherin failed to occur (Figure 2 S, V), ZO-1 was only detected in a few small foci (Figure 2 T, V) and, although a fibronectin matrix was present, it appeared less dense (Figure 2 W, X). An incipient nuclear alignment was sometimes observed (Supplementary Figure 3 K, arrowheads), but cells did not polarize their F-actin into apically enriched adhesion belts (compare Supplementary Figure 3 G-I with J-L). Epithelial tissues other than somites (e.g. ectoderm and neural tube) did not present significant alterations after incubation with Blebbistatin (Supplementary Figure 4). Altogether, these results point to an indispensable role for NM II activity for the MET underlying somite formation.

RockOut-treated explants also showed perturbations in somite formation, although to a lesser extent (Figure 1 E). When compared to control explants (Figure 3 A-D; Supplementary Figure 5 A-C), RockOut treatment resulted in incomplete somitic clefts, such that S0 shared the somitocoel with SI and sometimes also with SII (Fig 3 E-H, arrows, Supplementary Figure 5 D-F, arrows). In contrast to the accumulation of fibronectin in the nascent clefts in controls (Figure 3 B, arrow), no fibronectin was observed in the incipient somitic clefts of RockOut-treated explants (Figure 3 F, arrows). These results suggest that ROCK I/II activity in the S-IV and S-III regions of the PSM (regions e and d in Figure 1 A) is required for the formation of individualized somites. In contrast, when the rostral-most PSM (stage S-I before culture, region b in Figure 1 A) was exposed to RockOut for 6 hours, it was indistinguishable from control explants, showing apical accumulation of ZO-1 (Figure 3 I, M, arrowhead) and N-cadherin (Supplementary Figure 5, G, J), nuclear alignment (Figure 3, K, O, Supplementary Figure 5, H, K) and a complete, fibronectin matrix-containing cleft (Figure 3 J, N). This indicates that ROCK I/II activity is not required for S-I to develop into a somite.

**Figure 3.**
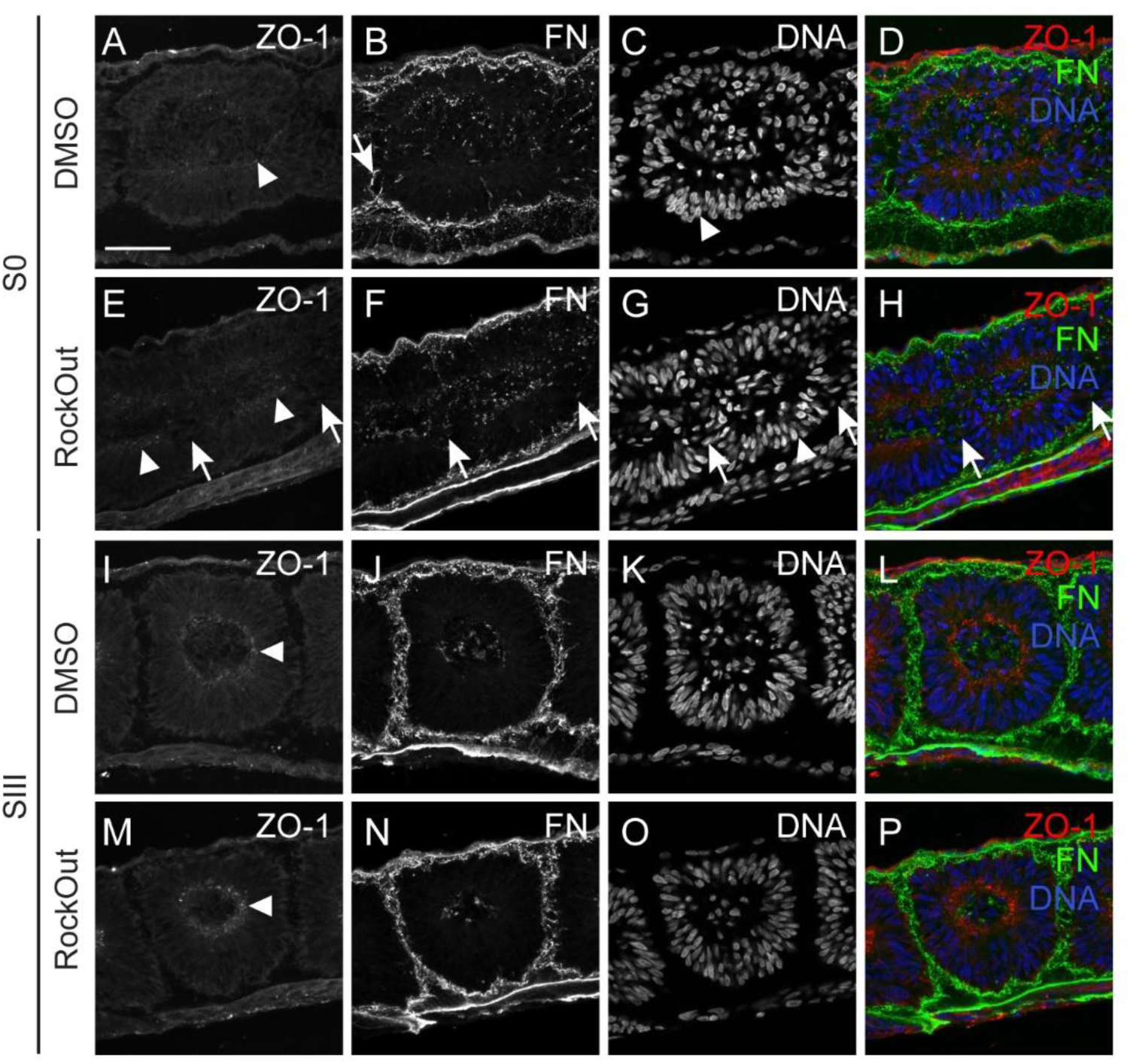
ROCK I/II inhibition impairs morphological somite formation, leading to deficient ZO-1 polarization and fibronectin assembly. (**A-P**)Sagittal sections of explants cultured in control (DMSO) medium (A-D, I-L) and their contralateral RockOut-treated halves (E-H, M-P) at S0 (A-H; region e in Figure 1 A) and SIII (I-P; region b in Figure 1 A) levels, immunostained for ZO-1 (first column), fibronectin (second column), stained for DNA (third column) and the respective merged image (fourth column). Explants were cultured for 6 hours. S0 in control explants show normal accumulation of ZO-1 (A, arrowhead), fibronectin assembly in the nascent cleft (B, arrow) and nuclear alignment (C, arrowhead). In contrast, S0 in contralateral RockOut-treated explants fails to form a clear cleft (E-H, arrows), although ZO-1 is generally polarized (E, arrowhead) and nuclei are aligned (G, arrowhead). At SIII level, both explants show normal ZO-1 polarization (I, M, arrowheads), fibronectin assembly (J, N) and nuclear alignment (K, O). Rostral on the left and dorsal on top. FN – fibronectin. ZO-1 – *Zonula occludens* protein 1. Scale bar: 50 µm.

Altogether, our data indicate that intracellular actomyosin contractility plays a role in periodic somite cleft formation, and that ROCK I/II-independent NM II activity is essential for MET.

### Blocking integrin-fibronectin binding perturbs segmentation clock oscillations and somitic cleft formation

Our next aim was to address the requirement for the fibronectin ECM surrounding the PSM in regulating segmentation clock oscillations and somite formation. PSM cells bind to the RGD motif of fibronectin through the α5β1 integrin, an interaction that plays a crucial role during somitogenesis (Girós et al., 2011; Yang et al., 1993). αv integrins, which have been described to bind the RGD motif and partially compensate for the absence of the α5β1 integrin in the mouse (Yang et al., 1999), are not detected in the PSM of the chick embryo (Gomes de Almeida et al., 2016). We cultured embryo half explants in the presence of a linear RGD peptide, which competes with fibronectin for integrin binding (Huveneers et al., 2008; Pierschbacher and Ruoslahti, 1984), and compared them to contralateral control explants (Figure 4 A, B). RGD-treated explants formed ill-defined somite-like condensations, although in approximately the same number as the contralateral control (Figure 4 C). This was not due to cell death (Supplementary Figure 2 E-F). Concomitantly, RGD-treated explants displayed alterations in *hairy1* (Figure 4 D, a-c), *meso1* (Figure 4 D, d-f) and *dll1* expression patterns (Figure 4 D, g-i), evidencing that integrin-fibronectin interactions via RGD are required for proper segmentation clock oscillations, *meso1* positioning and timely downregulation of *dll1* in the anterior PSM.

**Figure 4.**
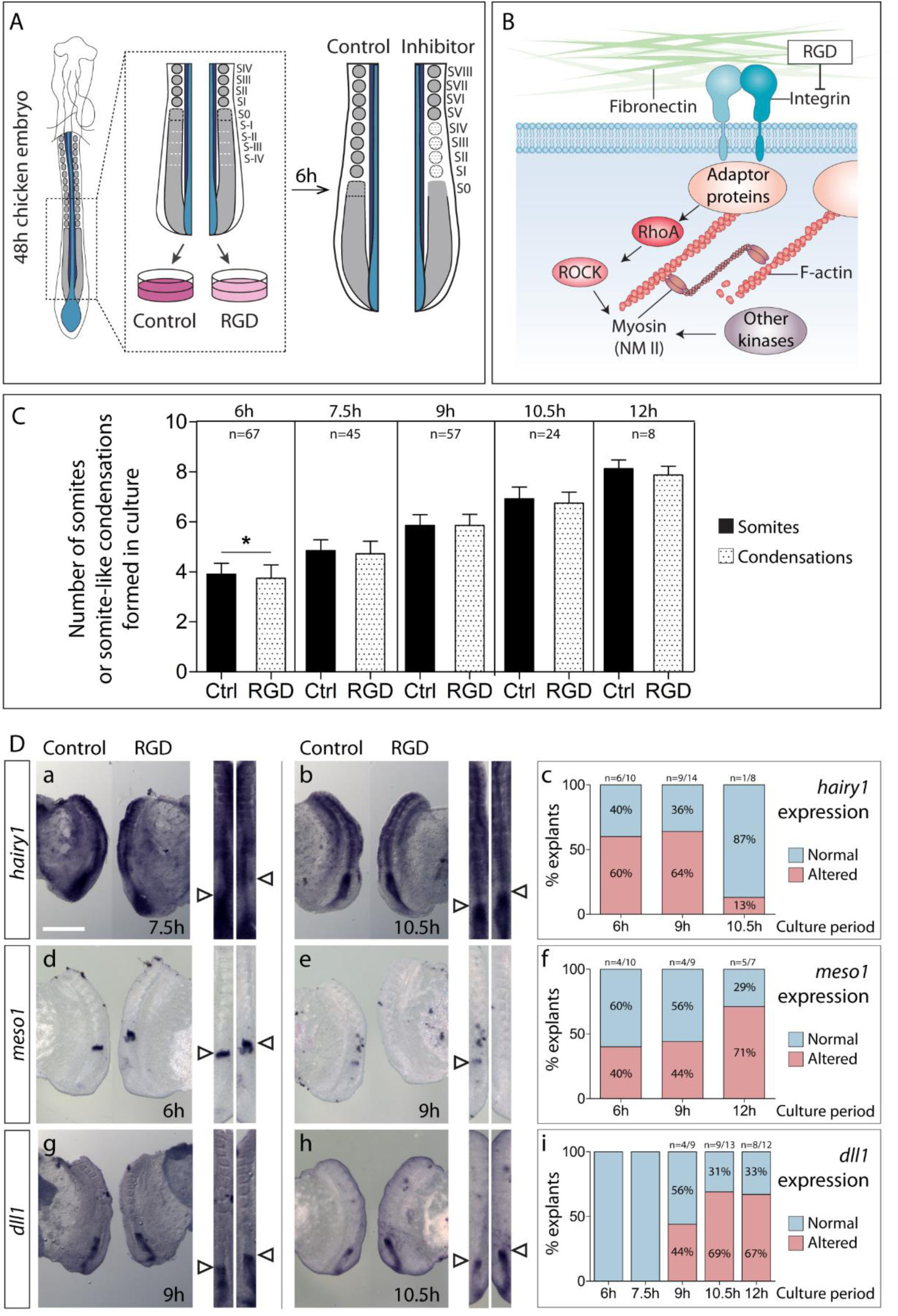
Integrin-fibronectin binding through RGD is required for timely *hairy1* and *meso1* expression and *dll1* downregulation. (**A**) Schematic representation of the explant culture system. Posterior explants of HH11-14 chick embryos were bisected along the midline and cultured for 6 to 12 hours. One side of the explant was cultured with RGD, while the contralateral half was cultured in control medium. **(B)** Schematic representation of the action of RGD. RGD competes with the RGD-binding pockets of integrins, interfering with their binding to the ECM. **(C)** Number of somites (black bars) or somite-like condensations (dotted bars) formed in culture in control and RGD-treated explants. p values were calculated using a paired Student’s t-test. *p<0.05. **(D)** Expression of *hairy1* (a, b), *meso1* (d, e) and *dll1* (g, h) in RGD-treated explants and contralateral controls at representative timepoints of culture. Straightened images of respective explant pairs (right) aligned by SIV. Rostral is on top. Percentage of RGD-treated explants with altered *hairy1*, *meso1* and *dll1* expression compared to the contralateral controls is shown in c, f and i, respectively. Impairing integrin-fibronectin binding with RGD alters *hairy1* and *meso1* expression relative to contralateral controls at 6 hours of culture onwards (a-f, arrowheads). *dll1* expression was altered at 9 hours of culture onwards (g-i, arrowheads). Scale bar: 500 µm.

When compared to the contralateral control, the area corresponding to S0 after 6 hours of culture with RGD (region e in Figure 1 A) showed deficient nuclear alignment (Figure 5 B, E, H, K, arrowheads) and N-cadherin polarization (Figure 5 A, G, arrowheads), accompanied by deficient fibronectin assembly in the nascent cleft (Figure 5 D, J arrowheads). At the level of SII (region c in Figure 1 A), complete somite individualization was impaired in RGD-treated explants (Figure 5 N, Q, T, W, arrowheads) and, although N-cadherin polarization appeared normal (Figure 5 M, S, arrowheads), cleft formation (Figure 5 Q, W, arrowheads; R, X) and fibronectin assembly between adjacent somites was deficient (Figure 5 P, V, arrowheads; R, X).

**Figure 5.**
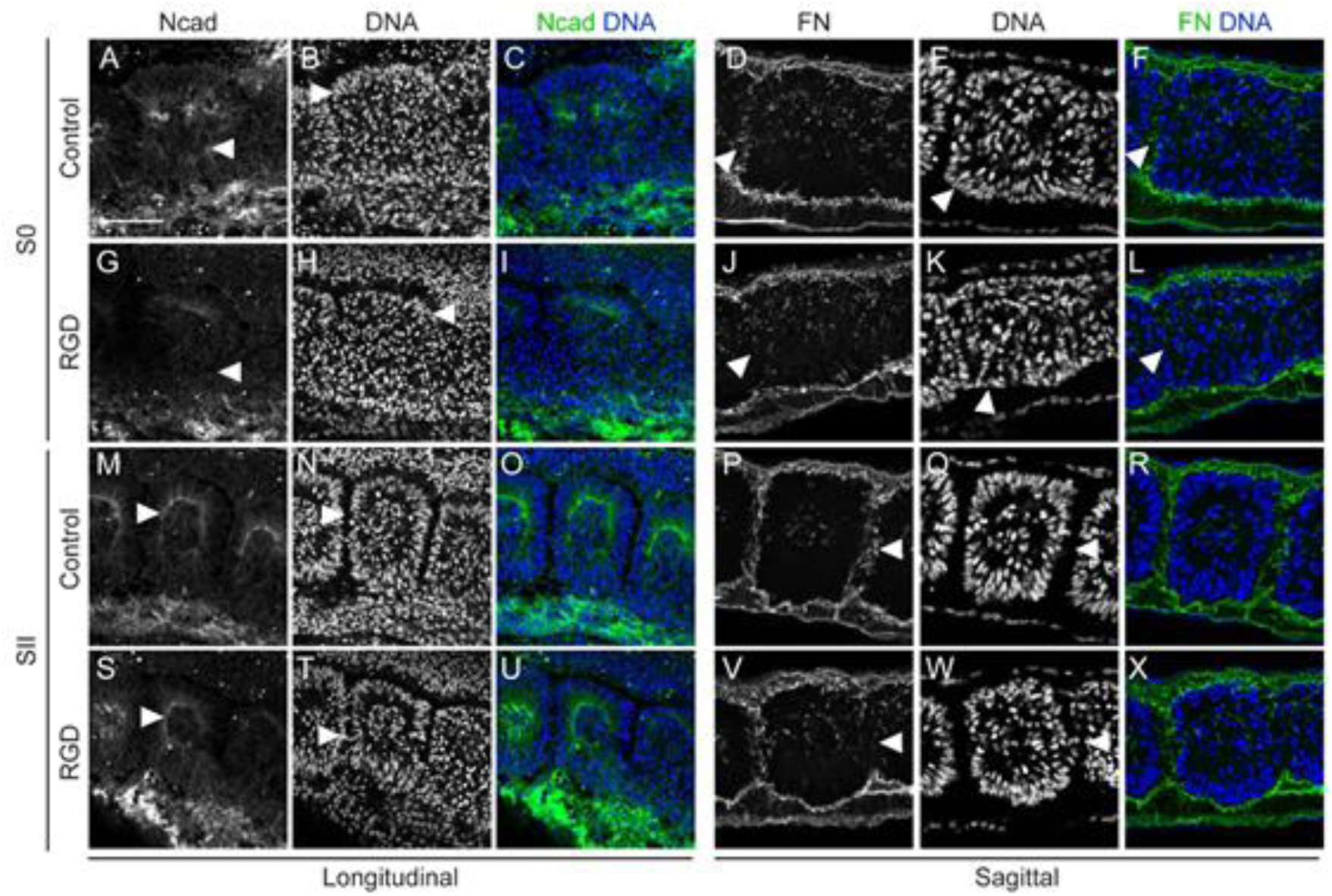
Inhibition of integrin-fibronectin binding impairs somitic cleft formation. (**A-X**) Longitudinal (left) and sagittal (right) sections of explants cultured in control medium (A-F, M-R) and their contralateral RGD-treated halves (G-L, S-X) at S0 (A-L; region e in Figure 1 A) and SII (M-X; region c in Figure 1 A) levels, immunostained for N-cadherin (first column), fibronectin (fourth column) and stained for DNA (second and fifth columns). Third and sixth columns show the respective merged images. Explants were cultured for 6 hours. At the S0 level, control explants show normal apical accumulation of N-cadherin (A, arrowheads) and nuclear alignment (B, E, arrowheads) in peripheral somitic cells as well as fibronectin assembly in the nascent cleft (D, F, arrowheads). All of these are deficient in their RGD-treated contralateral halves (G-L, arrowheads). At the level of SII, N-cadherin accumulation and nuclear alignment occurs normally in both control and RGD-treated explants (M, N, S, T, arrowheads), but fibronectin assembly between adjacent somites (P, V, arrowheads) and cleft formation (Q, W, arrowheads) are deficient in RGD-treated explants compared to contralateral controls. Ncad – N-cadherin; FN – fibronectin. Scale bars: 50 µm.

These findings implicate cell-ECM interactions, mediated by integrin-fibronectin binding via the RGD motif, in temporal control of *hairy1* expression, correct positioning of *meso1* expression, downregulation of *dll1* in the anterior PSM and somite morphogenesis.

### Impaired fibronectin matrix assembly results in altered segmentation clock dynamics, *meso1* expression and defects in somite morphogenesis

Another way to assess the relevance of the extracellular fibronectin matrix on the events leading up to somite formation is to perturb fibronectin assembly in the PSM and somites. To this end, primitive streak-stage embryos were electroporated with a construct expressing the 70kDa fibronectin fragment, a dominant-negative inhibitor of fibronectin matrix assembly (Figure 6 A, B; McKeown-Longo and Mosher, 1985; Sato et al., 2017). 70kDa-electroporated embryos exhibited a disrupted fibronectin matrix, composed of thinner fibrils when compared to control pCAGGs-electroporated embryos (Supplementary Figure 6 A; also see Figure 7 A, B). These embryos displayed multiple morphological defects, including kinked neural tube and detached tissues as well as perturbations in somite morphogenesis, which are all reminiscent of phenotypes obtained in previous studies interfering with fibronectin matrix deposition and/or with fibronectin-integrin binding (Supplementary Figure 6 B, C; Drake et al., 1992; Drake and Little, 1991; George et al., 1993; Girós et al., 2011; Takahashi et al., 2007). Although the average number of somite-like structures formed in 70kDa-electroporated embryos was similar to the number of somites in controls (Figure 6 C), the former were ill-defined, often appearing fused or crammed (Supplementary Figure 6 B e-f, C), closely resembling the somite-like condensations formed in RockOut- and RGD-treated explants (Figures 3 and 5).

**Figure 6.**
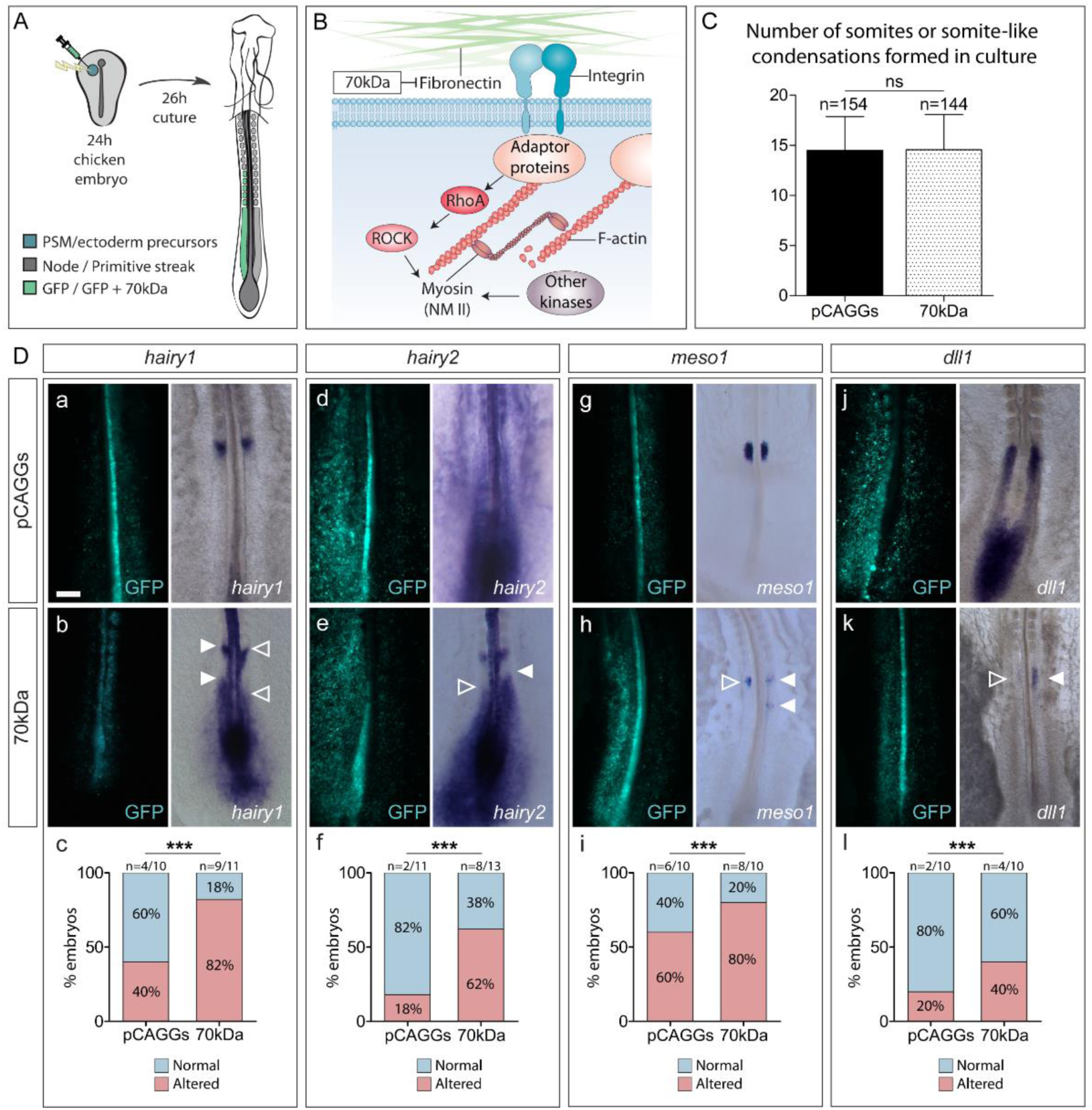
Impairing fibronectin assembly by electroporation with a 70kDa expressing vector perturbs segmentation clock oscillations, timely *meso1* expression and *dll1* downregulation. (**A**) Schematic representation of the electroporation strategy. PSM/ectoderm progenitors of primitive-streak stage embryos were electroporated with either a pCAGGs GFP-expressing vector (pCAGGs) alone, or co-electroporated with a 70kDa-expressing vector (70kDa) and were incubated for about 26 hours. **(B)** Schematic representation of the action of 70kDa. 70kDa disrupts the assembly of fibronectin matrix by competitively binding to the N-terminal self-assembly domains of the protein, impairing fibronectin fibril formation. **(C)** Number of somites (black bars) or somite-like condensations (dotted bars) formed in pCAGGs- and 70kDa-electroporated embryos after 26 hours. p values were calculated using a paired Student’s t-test. **(D)** Examples of the expression of *hairy1* (a-b), *hairy2* (d-e), *meso1* (g-h) and *dll1* (j-l) in pCAGGs-(top row) and 70kDa-electroporated embryos (middle row). Electroporated side is on right (a, b, g) or left (d, e, h, j, k) Perturbing the assembly of fibronectin on one side of the PSM leads to an asymmetric pattern of *hairy1* (b, arrowheads), *hairy2* (e, arrowheads), *meso1* (h, arrowheads) and *dll1* expression (k, arrowheads) in a higher percentage of embryos than in controls. Percentage of pCAGGs- and 70kDa-electroporated embryos with asymmetric expression of *hairy1*, *hairy2*, *meso1* and *dll1* between the electroporated PSM and the contralateral non-electroporated control PSM is shown in c, f, i and l, respectively. The number of embryos with an asymmetric pattern was significantly higher in 70kDa-electroporated embryos for all four genes studied. p values were calculated using a Chi-square test. *** p<0.01. Rostral is on top. Scale bar: 200 µm.

**Figure 7.**
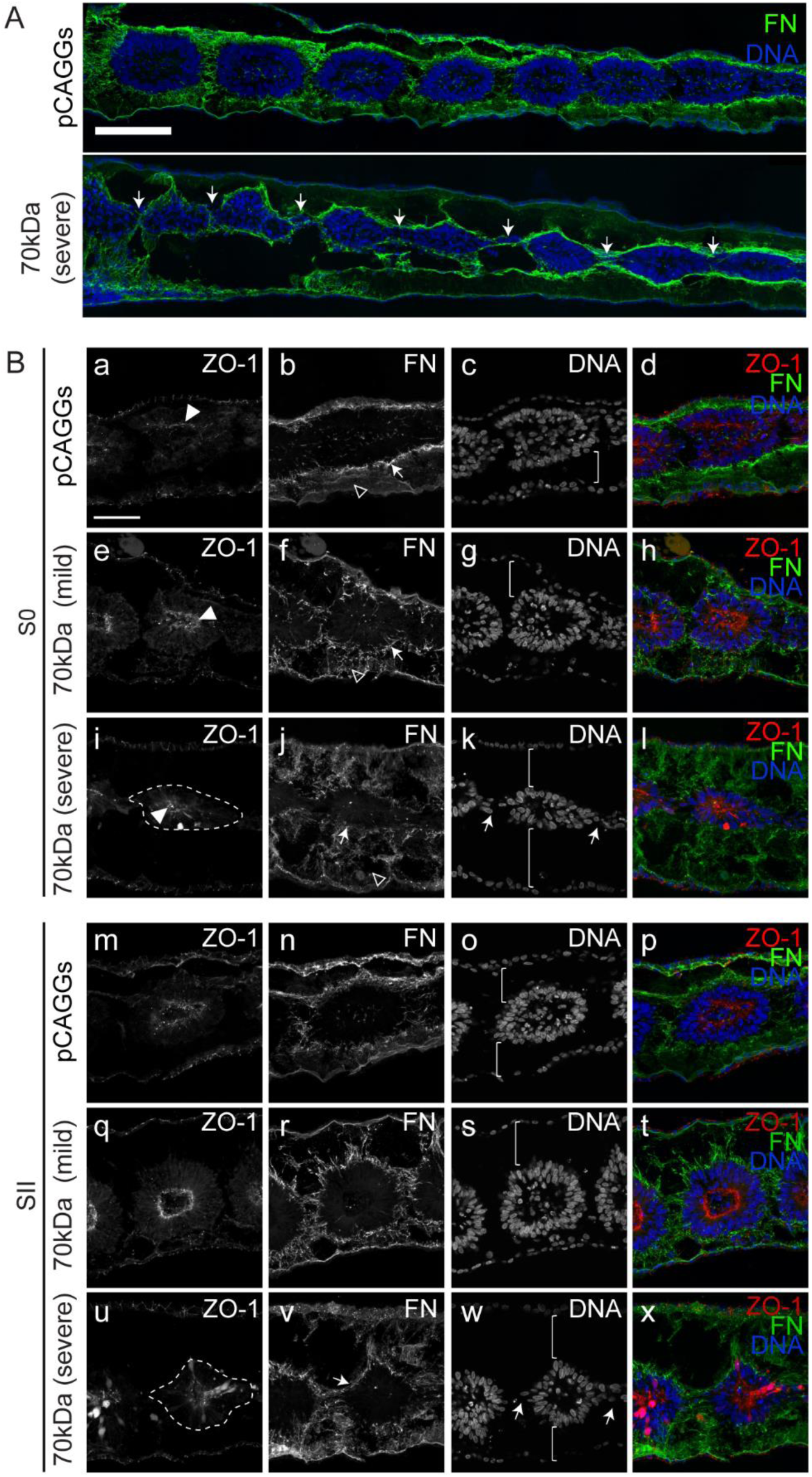
Somite morphology of 70kDa-electroporated embryos is severely compromised. (**A**) Sagittal sections of embryos electroporated with pCAGGs and 70kDa, the latter with a severe phenotype. Arrows point to deficient somitic clefts. Rostral to the left and dorsal on top. **(B)** Sagittal views of embryos electroporated with pCAGGs (a-d, m-p) and 70kDa (e-l, q-x) with either mild (e-h, q-t) or severe (i-l, u-x) phenotypes, at S0 (a-l) and SII (m-x) levels, immunostained for ZO-1 (first column) and fibronectin (second column) and stained for DNA (third column). Fourth column shows the merge of the respective channels. S0 of pCAGGs- and 70kDa-electroporated embryos all polarize ZO-1 normally (a, e, i, arrowheads) and this is maintained at SII levels (m, q, u), but the fibronectin matrix surrounding the somites of 70kDa-treated embryos is disrupted compared to pCAGGs-electroporated embryos (b, f, j, v, arrows). Somites of 70kDa-electroporated embryos are also severely detached from both the ectoderm and endoderm compared to embryos electroporated only with pCAGGS (third column, brackets), and the fibronectin matrix connecting the endoderm to the somites is severely compromised (second column, empty arrowheads). Somites of embryos electroporated with 70kDa with more severe defects also fail to fully detach from adjacent somites (k, w, arrows). Rostral to the left and dorsal to the top. Dashed lines indicate altered somite morphology. Experiments were performed in 6 (pCAGGs) and 7 (70kDa) biological replicates. FN – fibronectin. ZO-1 – *Zonula occludens* protein 1. Scale bars: 50 µm.

Next, we sought to evaluate the impact of inhibiting fibronectin matrix assembly on the molecular machinery underlying somite formation. Unilateral electroporation of the 70kDa fragment was performed (Figure 6 A) to allow the direct comparison of gene expression patterns within the same embryo. We observed a significant increase in the frequency of perturbations in the expression of embryonic clock genes *hairy1* (p<0.01) and *hairy2* (p<0.05), as well as in *meso1* (p<0.01) and *dll1* (p<0.01) expression, when compared with embryos electroporated with pCAGGs alone (Figure 6 D).

Consistent with these data, 70kDa-electroporated embryos have deficiencies in somite morphogenesis which, in severe cases, leads to incomplete somitic clefts (Figure 7 A, B k, w, arrows). Peripheral cells of nascent somites of 70kDa-electroporated embryos did, however, accumulate ZO-1 apically (Figure 7 B e, h, i, l) which was maintained as the somites matured (Figure 7 B q, u, t, x). Nevertheless, these somites were abnormal in shape and appeared smaller in severely affected embryos (Figure 7 B, i-I, u, x). In fact, the SI of the electroporated sides of 70kDa-treated embryos were significantly smaller in width than those of the contralateral control, while SV was significantly shorter in length (n=151; Supplementary Figure 7). In addition to defects in somite morphology, the ectoderm and endoderm were separated from the paraxial mesoderm in 70kDa-electroporated embryos, indicating that their fibronectin matrix was insufficient to hold these tissues together (brackets in Figure 7 B g, k, s, w).

We conclude that proper fibronectin matrix assembly in the PSM is required for timely clock gene expression dynamics, positioning of *meso1* expression and *dll1* downregulation in the rostral-most PSM, as well as for the complete separation and morphogenesis of somites.

## Discussion

### Fibronectin-dependent tissue mechanics coordinate segmentation clock dynamics and cleft formation

We have identified fibronectin-dependent tissue mechanics as a regulator of segmentation clock gene expression and the positioning of the presumptive somitic cleft in the chick embryo. Four independent treatments interfering with the mechanical properties of the PSM (Figure 8 A) consistently lead to asymmetric patterns of *hairy1* expression as well as incorrect positioning of *meso1* expression on the experimental versus the control sides of the same embryo. The similarity of the phenotypes obtained in experiments impairing fibronectin fibrillogenesis, cell-fibronectin interactions via RGD and blocking ROCK (Figure 8 B) suggests that the fibronectin matrix, the RGD-binding α5β1 integrin, and ROCK-dependent actomyosin contractility are part of the same pathway. The α5β1 integrin can transduce mechanical signals by ROCK activation (Schiller et al., 2013) and this requires the binding of α5β1 to two sites of fibronectin, namely the RGD and the synergy site (Friedland et al., 2009). Surprisingly, although the RGD site of fibronectin is crucial for somitogeneses (Girós et al., 2011), removing the synergy site of fibronectin (*Fn1^syn/syn^*) had no effect on mouse embryonic development (Benito-Jardón et al., 2017). However, cells expressing αv-integrins form strong adhesions on fibronectin lacking the synergy site and can compensate for the inability of α5β1 to mediate adhesion strengthening on this fibronectin (Benito-Jardón et al., 2017), and αv-integrins can partially compensate for the complete absence of α5β1 during mouse somitogenesis (Yang et al., 1999). Alternatively, ROCK can be activated indirectly, for example through cadherin engagement and subsequent adherens junction formation (Burute and Thery, 2012; Schwartz and DeSimone, 2008), which occurs upon fibronectin-induced polarization of peripheral PSM cells (Martins et al., 2009).

**Figure 8.**
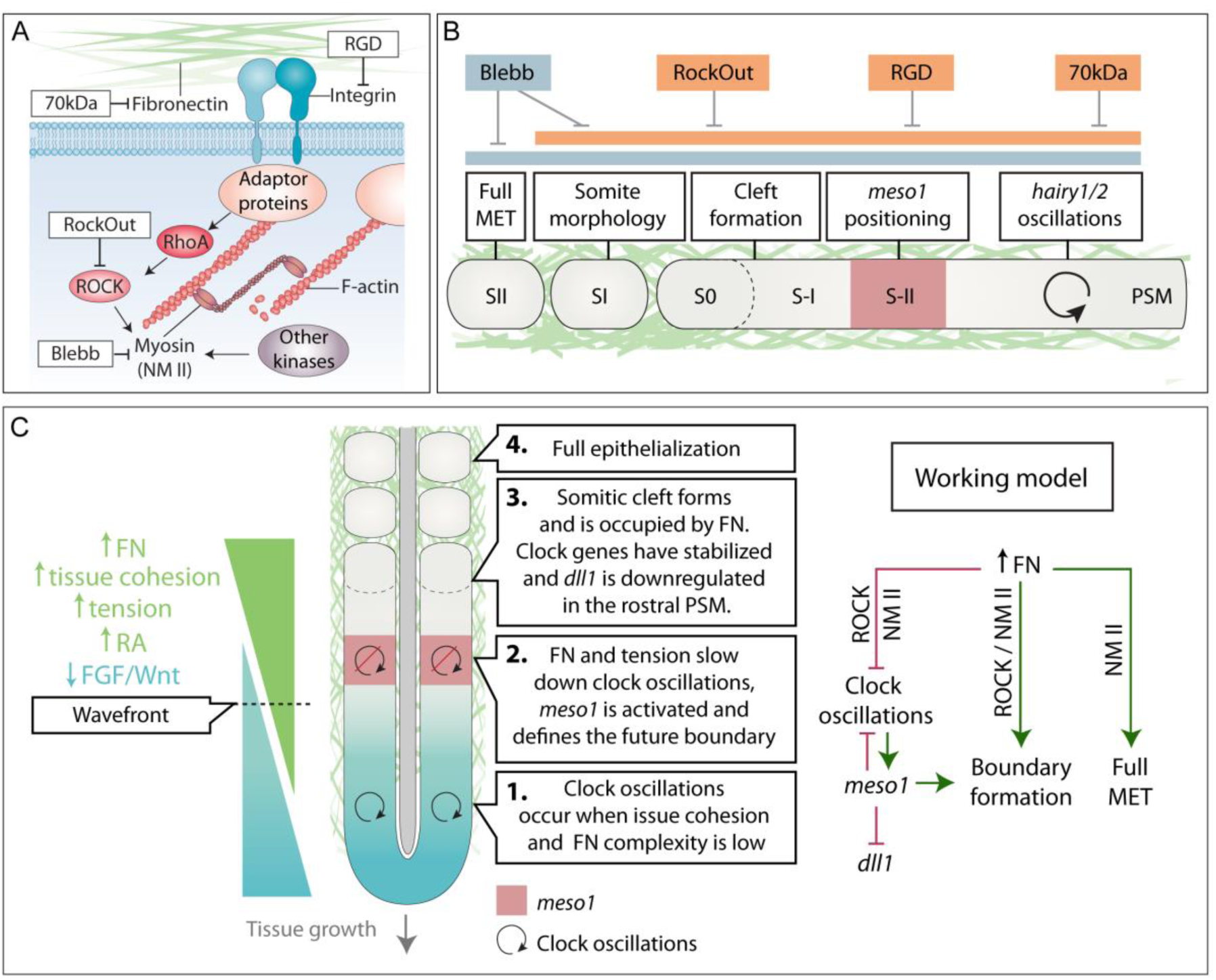
Working model illustrating a fibronectin-dependent mechanotransduction pathway in chick somite formation. (**A**) Schematic representation of the targets of experimental inhibition used in this study. **(B)** Effect of the different treatments on somitogenesis. Interfering with NM II (Blebb) or ROCK I/II (RockOut) activity, integrin-fibronectin binding (RGD), or fibronectin matrix assembly (70kDa), leads to altered segmentation clock dynamics, misregulated *meso1* expression, impaired or deficient cleft formation and alterations in somite morphology. Interfering directly with NM II further disrupts full somite epithelialization. **(C)** Working model proposing that an increase of fibronectin matrix complexity and tissue cohesion in the PSM, from caudal to rostral, is sensed by cells as a tensional gradient, with anterior PSM cells being in the stiffest environment. This increased tensional gradient occurs concomitantly with a decrease in Fgf/Wnt levels and an increase in RA. As cells reach the wavefront and sense a threshold of stiffness (due to fibronectin-dependent tissue cohesion), ROCK activity is increased and so is actomyosin contractility. This in turn slows down clock oscillations and stabilizes Notch signaling, which activates *meso1* expression in the S-II/S-I region, setting the stage for cleft formation and *dll1* downregulation in the rostral-most PSM later on. As the clefts form, somites bud off as SI. Finally, ROCK-independent NM II activity is required for full somite epithelialization into SII.

It is well established that Notch signaling plays a role in the segmentation clock and is also required for timely *meso1* activation (Saga, 2012). Hence, the mechanical environment may be regulating Notch signaling in the PSM. In agreement with our results, chicken embryos electroporated with RNAi constructs against integrin β1 showed alterations in *hairy2, lfng* and *meso1* expression in the PSM (Rallis et al., 2010). Mouse embryos where the fibronectin RGD site has been substituted with an RGE sequence (*Fn1^RGE/RGE^*) also showed asymmetric and/or dampened expression of *Lnfg* and *Hes7* in the PSM (Girós et al., 2011) and *EphA4*, a direct target of Mesp2 in the anterior PSM (Nakajima et al., 2006), was diffusely expressed or absent (Girós et al., 2011). Finally, combined roles of integrin α5β1 and Notch are required for zebrafish somitogenesis (Jülich et al., 2005).

Exactly how tissue mechanics regulate Notch signaling is unknown. It is becoming increasingly clear that mechanics play a crucial role in Notch activation (Gordon et al., 2015; Luca et al., 2017; Meloty-Kapella et al., 2012) and sustained Notch signaling in the *Drosophila* notum requires actomyosin contractility in both signal sending and receiving cells (Hunter et al., 2019). Intriguingly, clock oscillations in mouse tailbud PSM explants cultured on fibronectin are sustained in the presence of a ROCK inhibitor, suggesting that ROCK activity must be low for the maintenance of clock oscillations and that an increase in ROCK activity normally stops segmentation clock oscillations in this system (Hubaud et al., 2017). Moreover, in the same study, cell adhesion to fibronectin was linked to nuclear localization of Yes-associated protein (YAP), an intracellular sensor of cell mechanics, and dampening and eventual arrest of *Lnfg* oscillations was found to be YAP-dependent (Hubaud et al., 2017). ROCK-mediated actomyosin contractility is known to promote the nuclear localization of YAP in several cell types (Piccolo et al., 2014) and YAP-null mouse mutants (Morin-Kensicki et al., 2006) have a phenotype very similar to that of integrin α5-null mutants (Yang et al., 1993) and *Fn1^RGE/RGE^* embryos (Girós et al., 2011), suggesting that they contribute the same processes during early embryo development. Thus, it is conceivable that increased fibronectin-dependent tissue cohesion may translate into increased ROCK activity and actomyosin contractility, promoting sustained Notch signaling and nuclear localization of YAP, leading to the dampening and eventual arrest of clock oscillations. Further studies are needed to test this hypothesis.

Altogether, our results show that perturbation of the normal PSM tissue mechanics leads to a dysregulation of segmentation clock oscillations and the mispositioning of the segmental border, indicating that the mechanical properties of the PSM modulate Notch signaling and coordinate the translation of clock oscillations into periodic segmental border formation.

### Somite cleft formation and cell epithelialization have different mechanical requirements

In the rostral PSM, Mesp2/Meso1 activates the expression of EphA4, which interacts with EphrinB2 in cells rostral to the presumptive cleft, causing cell-cell repulsion and the formation of an incipient cleft (Nakajima et al., 2006; Watanabe et al., 2009). Then, fibronectin matrix assembly within the cleft stabilizes it (Jülich et al., 2015; Rifes and Thorsteinsdóttir, 2012) and promotes the epithelialization of cells rostral to the cleft (Martins et al., 2009). This can be defined as the first step of morphological somite individualization. The second step is defined as the complete epithelialization of the remaining cells of the nascent somite and lasts until SII, when all somitic cells have acquired a spindle-like shape and are organized into a rosette (Martins et al., 2009).

RockOut-, RGD- and 70kDa-treated embryos (Figure 8 A, B) all show perturbations in segmentation clock gene expression, abnormal positioning of *meso1* and defects in the somitic clefts. Although somite morphology is also perturbed, the acquisition of the spindle-shape cell morphology which occurs as S0 develops into SII does not appear to be significantly perturbed. Thus, the first step of morphological somite formation is affected but the second step is not. In contrast, Blebbistatin-treated explants not only have the defects listed above, but cells that had already upregulated *meso1* before the addition of the drug and formed an incipient cleft during culture, were completely unable to epithelialize. In fact, Blebbistatin-treated explants formed only 1 or 2 somites, and their cells did not acquire the elongated, spindle-shape typical of SI and SII somites. Hence, Blebbistatin affects both steps of morphological somite formation.

RockOut targets NM II activity indirectly by inhibiting ROCK I and II, two of the kinases that activate NM II (Newell-Litwa et al., 2015). In contrast, Blebbistatin directly targets the NM II ATPase. Our results thus raise the possibility that the acquisition of the spindle-shaped morphology of cells may be dependent on another NM II activator. Interestingly, Ca^++^/calmodulin signaling can activate NM II and inhibiting calmodulin was shown to block the acquisition of this morphology during chick somitogenesis (Chernoff and Hilfer, 1982).

### A gradient of fibronectin-dependent tissue mechanics as a player in the PSM wavefront of maturation

While the waves of Notch oscillatory activity travel through the entire length of the PSM, they are only translated into segments in the anterior-most region of the tissue. Opposing gradients of Fgf/Wnt and Retinoic acid (RA) in the PSM are thought to define the wavefront, which marks the region where PSM cells become competent for somite formation (Hubaud and Pourquié, 2014). Rostral to the wavefront, segmentation clock oscillations progressively slow down until they reach a halt and cells become part of a somite (Palmeirim et al., 1997).

Herein, we propose a model (Figure 8 C) where a posterior to anterior gradient of fibronectin-dependent tissue cohesion is interpreted by the PSM cells as an increasing gradient of mechanical tension (Figure 8 C). We propose that the fibronectin-dependent tensional state of the PSM at the level of the wavefront acts as a threshold that activates a mechanotransduction signaling cascade, ensuring the correct spatio-temporal conversion of the cyclic expression of segmentation clock genes into periodic *meso1* expression, which in turn defines the next somitic cleft (Figure 8 C). In support of this hypothesis, a gradient of increasing paraxial mesoderm stiffness from the tail to rostral somites has been identified in the chick embryo (Bénazéraf et al., 2017; Marrese et al., 2019). During its maturation, the PSM thus integrates a combination of chemical and mechanical signals, namely gradients of Fgf/Wnt and RA (Aulehla and Pourquié, 2010) and, simultaneously, a progressive increase in fibronectin-dependent tissue cohesion (Figure 8 C). Fibronectin matrix-dependent tissue mechanics would thus be a key contributor to the PSM wavefront of maturation in that it regulates where and when the next somitic cleft is positioned in the anterior PSM (Figure 8 C). Hence, we propose that a mechanotransduction pathway downstream of fibronectin plays a major role in the translation of cyclic waves of expression of segmentation clock genes into the periodic morphogenesis of somites.

## Materials and Methods

### Embryos

Fertilized chicken (*Gallus gallus)* eggs were obtained from commercial sources (Sociedade Agrícola Quinta da Freiria or Pintobar Exploração Avícola, Lda, Portugal) and incubated at 37.5°C in a humidified chamber until the desired HH stage (HH4 or HH11-14; Hamburger and Hamilton, 1992). Somite nomenclature is according to Pourquié and Tam (2001).

### Embryo explant culture and chemical treatments

Explant tissues of HH11-14 embryos were collected and cultured as previously described (Palmeirim et al., 1997; Rifes et al., 2007). Embryos were bisected along the midline and then cut transversally rostral to somites IV and Hensen’s node. The two contralateral halves thus retained half of the neural tube and notochord as well as the first four somites and the PSM, with all remaining neighboring tissues intact. Explants were placed on top of a polycarbonate filter floating on M199 medium supplemented with 10% chick serum, 5% fetal calf serum and 100 U/ml of penicillin and streptomycin (Palmeirim et al., 1997). Explants were then cultured at 37°C with 5% CO2 from 6 to 12 hours.

InSolution™Blebbistatin (Calbiochem) and RockOut (Calbiochem) diluted in DMSO were used at a final concentration of 50 µM in culture medium. Equal volumes of DMSO (Sigma-Aldrich) were used as control for both drugs. The linear RGD peptide (GRGDS - G4391, Sigma) was diluted in culture medium and used at 0.9 mM, while control explants were cultured in medium only. RGD peptide efficiency was confirmed in a cell adhesion assay (Danen et al., 2002; Pierschbacher and Ruoslahti, 1984) before using it on explants.

### Embryo electroporation and *ex ovo* culture

HH4-5 embryos were electroporated on one (randomly selected) side of the primitive streak in the presumptive PSM and/or ectoderm and cultured *ex ovo* using the Early Chick culture method (Chapman et al., 2001). The electroporation mixture contained plasmid DNA at 0.5-1 µg/µl mixed with 0.4% Fast Green for visualization. Embryos were submerged in an electroporation chamber filled with Tyrode’s saline and three pulses of 6-9 V, 50 ms each, at 350 ms intervals were applied. Control embryos were electroporated with pCAGGs containing a GFP reporter (pCAGGs-GFP; abbreviated pCAGGs). pCAGGs-70kDa qFN1 was kindly provided by Yuki Sato (Sato et al., 2017) and was co-electroporated with the pCAGGs-GFP plasmid in experimental embryos (treatment abbreviated 70kDa). Electroporated embryos were screened for GFP after fixation to select embryos with an intense signal on only one side to process for whole mount morphological analysis, *in situ* hybridization experiments and transverse sectioning. For morphological analysis in sagittal sections, the embryo side electroporated with pCAAGGs was compared to the 70kDa-electroporated side of other same stage embryos.

### Cryosectioning and immunohistochemistry

Cryosectioning was performed on embryo explants and whole embryos fixed in 4% paraformaldehyde in 0.12 M phosphate buffer containing 4% sucrose and processed for cryoembedding. Fixed samples were washed in 0.12 M phosphate buffer with 4% and 15% sucrose and then embedded in 7.5% gelatin in 0.12 M phosphate buffer containing 15% sucrose, frozen on dry ice-chilled isopentane and stored at −80°C until sectioning. Cryostat sections (10-30 µm) were processed for immunofluorescence as previously described (Gomes de Almeida et al., 2016). Permeabilization of sections was performed with 0.2% Triton-X100 in phosphate buffered saline (PBS). 5% bovine serum albumen (BSA) or a combination of 1% BSA and 10% Normal Goat Serum (NGS) in PBS were used for blocking depending on the presence or absence of anti-fibronectin antibodies, respectively. Primary and secondary antibodies were diluted in 1% BSA in PBS. Sections were incubated with primary antibodies overnight at 4°C and with secondary antibodies for 1 hour at room temperature. For whole-mount immunodetection, explants were fixed in 4% paraformaldehyde in PBS and processed as previously described (Martins et al., 2009; Rifes and Thorsteinsdóttir, 2012). 1% Triton-X100 in PBS was used for permeabilization and 1% BSA in PBS was used for blocking and antibody dilution. Antibody incubation was performed overnight at 4°C.

The following primary antibodies were used: anti-ZO-1 (Zymed, #40-2200, 1:100 or Invitrogen, #33-9100, 1:100); anti-N-cadherin (BD Biosciences, #610920, 1:100); anti-fibronectin (Sigma, #F-3648, 1:400), anti-activated caspase3 (Cell Signaling, #9661, 1:1000) and anti-GFP (Invitrogen, #A11122, 1:100). For F-actin staining we used Alexa 488-conjugated phalloidin (Invitrogen, 1:40) and for staining DNA we used ToPro3 (Invitrogen, 1:500) in conjunction with ribonuclease A (Sigma, 10 µg/ml), 4% Methyl Green (Sigma, diluted 1:250; Prieto et al., 2015) or 4′,6-diamidino-2-phenylindole (DAPI, 5µg/ml in PBS with 0.1% Triton-X100). For detection of the primary antibodies the adequate secondary goat anti-mouse and anti-rabbit Alexa 488-, Alexa 568-or Alexa 546-conjugated F’ab fragments from Invitrogen were used (#A-11017, #A-21069, #A-11071, #A-11019, #A-11070, 1:1000). Immunohistochemistry was performed on at least 6 different explants/embryos and the respective controls for each treatment (Blebbistatin n=13; RockOut n=15; RGD n=13; 70kDa n=7/pCAGGs n=6).

### *In situ* hybridization

*In situ* hybridization using DIG-labeled RNA probes was performed as described previously (Henrique et al., 1995) with minor alterations (Gomes de Almeida et al., 2016). RNA probes were synthetized from linearized plasmids: *dll1* (Henrique et al., 1995), *meso1* (Buchberger et al., 1998), *hairy1* (Palmeirim et al., 1997) and hairy2 (Jouve et al., 2000).

### Statistical analysis

Paired Student’s t-tests were performed to assess for differences in the number of somites formed in Blebbistatin-, RockOut- and RGD-treated explants relative to the respective controls, and in embryos electroporated with pCAGGs only and pCAGGs + 70kDa. Differences in the frequency of morphological and gene expression phenotypes found in 70kDa-electroporated embryos compared to pCAGGs-electroporated control embryos was tested through a Chi-square test. Differences in somite size between pCAGGs- or 70kDa-electroporated sides compared to the control (non-electroporated) sides of embryos was tested through a nested ANOVA. The side in which the embryo was electroporated (left or right) was nested in the treatment (non-electroporated, pCAGGs-electroporated or 70kDa-electroporated) to account for a potential variability between the two sides. Statistical significance was set at p<0.05. Statistical analyses were performed in Statistica 10 (https://statistica.software.informer.com/10.0/), Graphpad Prism 5 (https://graphpad-prism.software.informer.com/5.0/) and RStudio (https://rstudio.com/).

### Sample preparation and imaging

Whole mount explants were gradually dehydrated in methanol and cleared in methylsalicylate (Sigma-Aldrich) as described previously (Martins et al., 2009; Rifes and Thorsteinsdóttir, 2012), except for phalloidin-labelled embryos and explants, where a shorter series of ethanol dehydration series was used. Cryostat sections were mounted in Vectashield (Vector Laboratories) or in 5mg/ml propyl gallate in glycerol/PBS (9:1) with 0.01% azide. Immunofluorescence images were taken on a confocal Leica SPE microscope, following imaging acquisition steps described previously (Rifes and Thorsteinsdóttir, 2012). Imaging of electroporated embryos and explants processed for *in situ* hybridization was performed using a Zeiss LUMAR V12 Stereoscope coupled to a Zeiss Axiocam 503 color 3MP camera. Image analysis was performed using Fiji v. 1.49 (https://imagej.net/Fiji) software. Image histogram corrections and, when appropriate, maximum intensity projections of immunofluorescence confocal stacks were produced in Fiji and exported as TIFF files. When applicable, contiguous images were stitched together into a single image using the pairwise stitching Fiji plugin (Preibisch et al., 2009). For the analysis of in situ hybridization patterns along the PSM explants, the Fiji plugin Straighten (Kocsis et al., 1991) was used.

## Acknowledgements

We thank Dr. Yuki Sato for generously sharing the pCAGGs-q70kDa construct and Inês Fragata for help with the statistical analysis. This work was supported by Fundação para a Ciência e a Tecnologia (FCT, Portugal) projects PTDC/SAU-OBD/103771/2008, PTDC/BEXBID/5410/2014, UID/BIA/00329/2013, UID/BIM/04773/2019 CBMR, and FCT scholarships SFRH/BD/86980/2012 (PGA) and SFRH/BD/37423/2007 (PR). Imaging and image analysis were done in the Microscopy Facility at the Faculty of Sciences of the University of Lisbon and the Light Microscopy Unit of CBMR-UAlg, nodes of the Portuguese Platform for BioImage (reference PPBI-POCI-01-0145-FEDER-022122). Finally, we are grateful to Isabel Palmeirim and all members of our laboratories for their support and helpful discussions.

## Competing interests

The authors declare no competing or financial interests.

## Author contributions

Conceptualization: P.G.A., P.R., R.P.A., S.T.; Methodology: P.G.A., P.R., R.P.A., S.T.; Validation: P.G.A, P.R., A.P.M.J., G.G.P., R.P.A., S.T.; Formal analysis: P.G.A.; Investigation: P.G.A., P.R., A.P.M.J., G.G.P.; Resources: R.P.A., S.T.; Writing - original draft: P.G.A., S.T.; Writing - review & editing: P.G.A, P.R., A.P.M.J., G.G.P., R.P.A., S.T.; Visualization: P.G.A., P.R., R.P.A., S.T.; Supervision: R.P.A., S.T.; Project administration: P.G.A., R.P.A., S.T.; Funding acquisition: R.P.A., S.T.

---

**Supplementary Figure 1.**
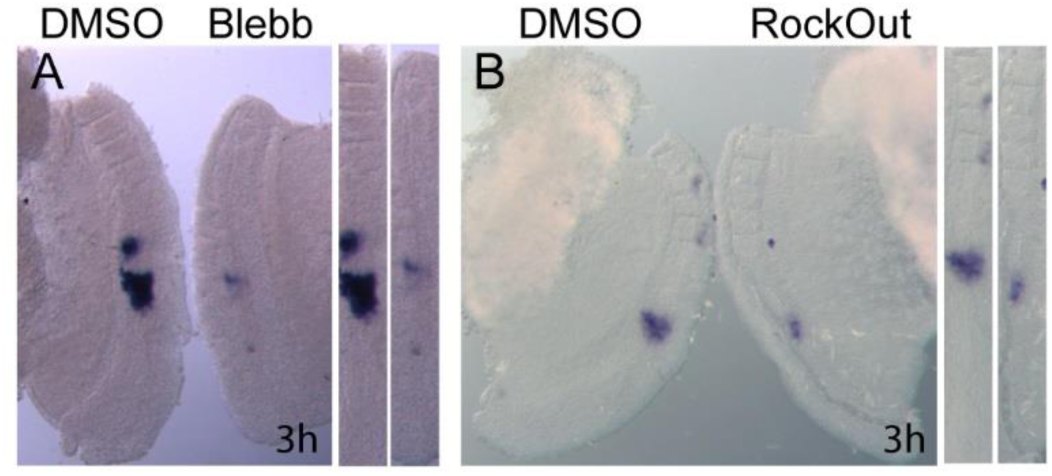
Blocking NM II or ROCK I/II activity for 3 hours leads to altered *meso1* expression. (**A**, **B**) m*eso1* expression evaluated by *in situ* hybridization in explants treated for 3 hours with Blebbistatin (**A**) or RockOut (**B**) and in the respective contralateral controls, shows that *meso1* expression is already altered two clock cycles after adding the drugs. Straightened images of the respective explant pairs (right) were aligned by SIV. Rostral is on top.

**Supplementary Figure 2.**
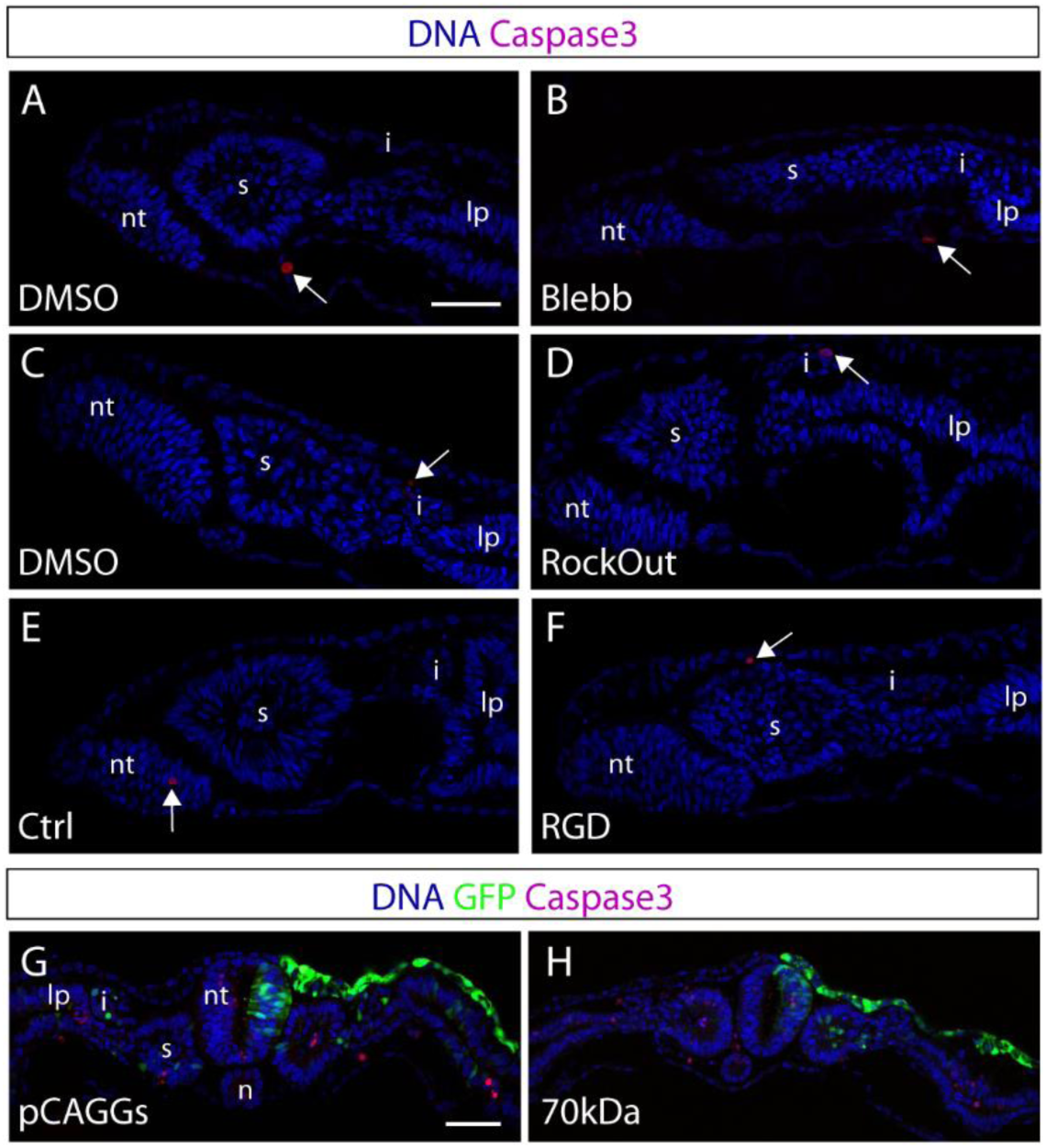
Apoptosis is not enhanced in the experimental culture conditions used. (**A**, **H**) Immunostaining for activated Caspase3 in transverse sections of control and contralateral explants treated with either Blebbistatin (**A, B**), RockOut (**C, D**), or RGD (**E, F**) for 6 hours, and in pCAGGs- and 70kDa- electroporated embryos (**G, H**). DNA (blue), activated Caspase3 (magenta) and GFP (green). Arrows indicate Caspase3-positive cells. Apoptosis levels are not increased by any of the treatments. Blebb – Blebbistatin; nt – neural tube; s – somite; i – intermediate mesoderm; lp – lateral plate mesoderm. Dorsal is on top. Scale bars: 50 µm.

**Supplementary Figure 3.**
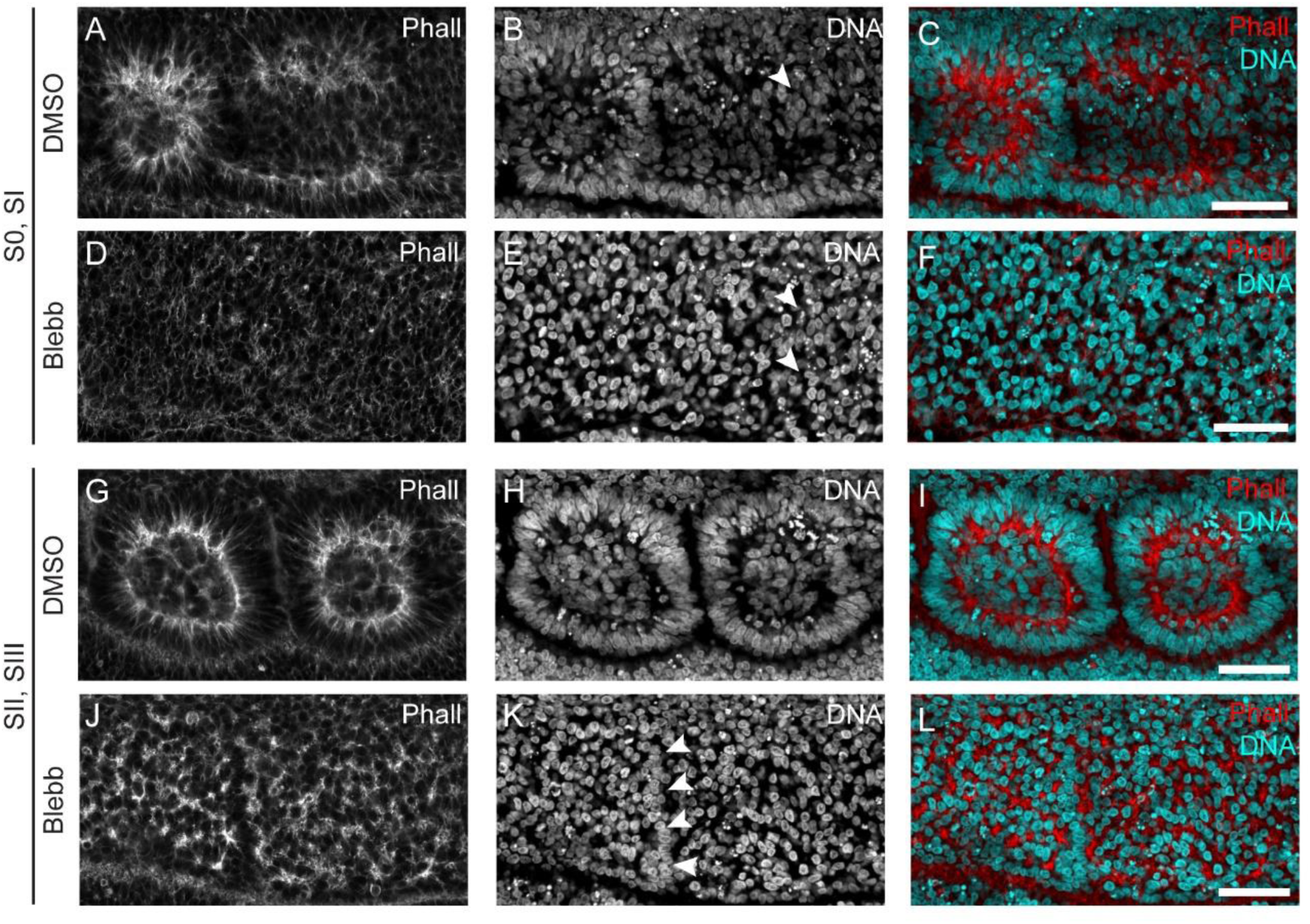
NM II inhibition impairs F-actin apical enrichment and nuclear alignment in somites formed during culture. (**A**, **L**) Sagittal optical sections of DMSO-treated explants (A-C, G-I) and contralateral Blebbistatin-treated explants (D-F, J-L) stained for F-actin and DNA. Rostral PSM epithelialization of DMSO-treated explants occurs normally, with F-actin apical enrichment (A) and nuclear alignment (B) in S0 and SI. At the same axial level in the contralateral Blebbistatin-treated explant (D-F), F-actin staining is dispersed (D) and nuclei do not align (E). SII and SIII of DMSO-treated explants are epithelial (G-I), composed of an outer cell layer with aligned nuclei (H) and elongated cells with apically enriched F-actin (G). At the equivalent axial level in the Blebbistatin-treated explant (J-L), the somitic segments were severely affected. There is only a slight nuclear alignment at the prospective inter-somitic border (K, arrowheads) and F-actin aggregates into dispersed and separate foci (J, L). Rostral to the left and dorsal on top. Blebb – Blebbistatin; Phall – phalloidin F-actin staining. Scale bars: 50 µm.

**Supplementary Figure 4.**
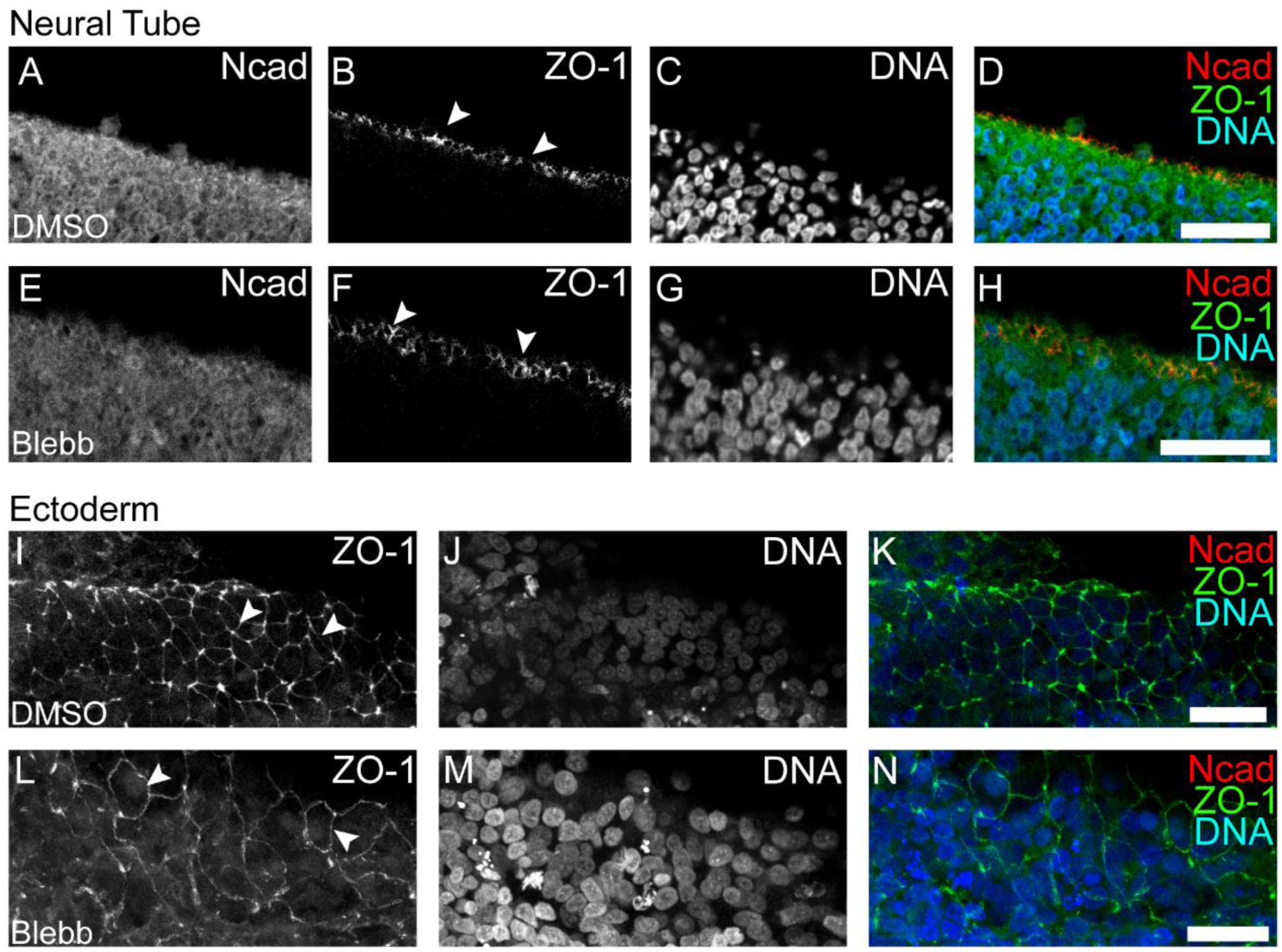
Epithelial structure of the neural tube and ectoderm are unaffected by NM II inhibition. (**A**, **N**) Control explants (A-D and I-K) showed strong apically located ZO-1 labeling (B, I, arrowheads) both in the neural tube (A-D; transverse optical section, medial on top) and in the overlying surface ectoderm (I-K; dorsal view of ectoderm). In the presence of Blebbistatin (E-H, L-N), ZO-1 labeling remained restricted to the apical end of neural tube cells (F, arrowheads) and of surface ectoderm cells (L, arrowheads). Blebb – Blebbistatin; Ncad – N-cadherin. ZO-1 – *Zonula occludens* protein 1. Arrowheads point to ZO-1 labeling. Scale bars: 20 µm.

**Supplementary Figure 5.**
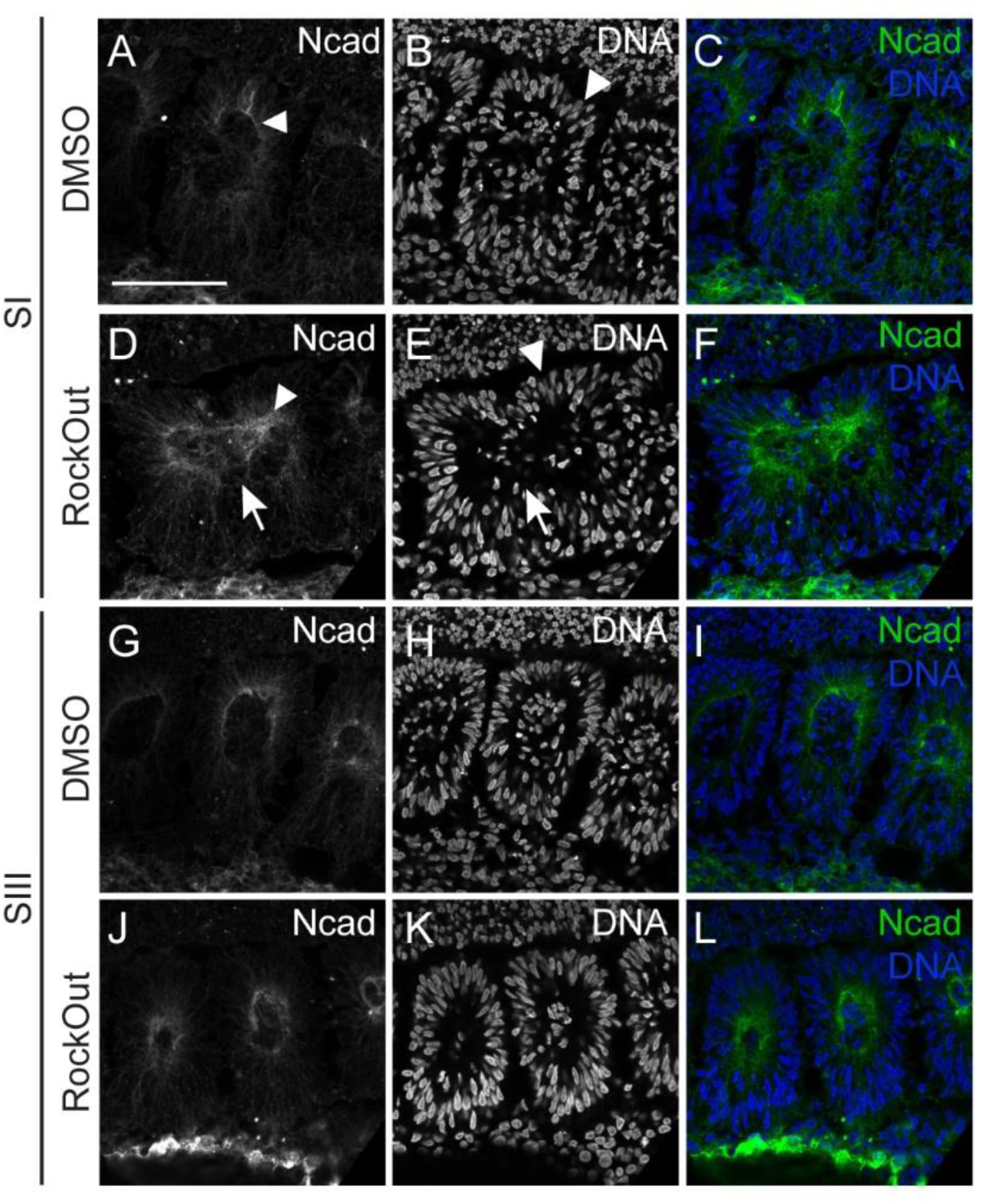
ROCK I/II inhibition impairs apical polarization of N-cadherin in nascent somites. (**A**, **L**) Longitudinal sections of explants cultured for 6 hours in control (DMSO) medium (A-C, G-I) and their contralateral RockOut-treated halves (D-F, J-L) at SI (A-F) and SIII (G-L) levels, immunostained for N-cadherin (first column) and stained for DNA (second column). Third column shows the respective merge of all stainings. SI in control explants shows normal accumulation of N-cadherin (A, arrowhead) and nuclear alignment (B, arrowhead). In contrast, SI in contralateral RockOut-treated explants fail to form a clear cleft (D-F, arrows) leading to partially fused somites. N-cadherin is, however, partially polarized (D, arrowhead) and nuclei are aligned (E, arrowhead). At SIII level, both explants show normal N-cadherin polarization (G, J) and nuclear alignment (H, L). Rostral on the left and midline on top. Ncad - N-cadherin. Scale bars: 50 µm.

**Supplementary Figure 6.**
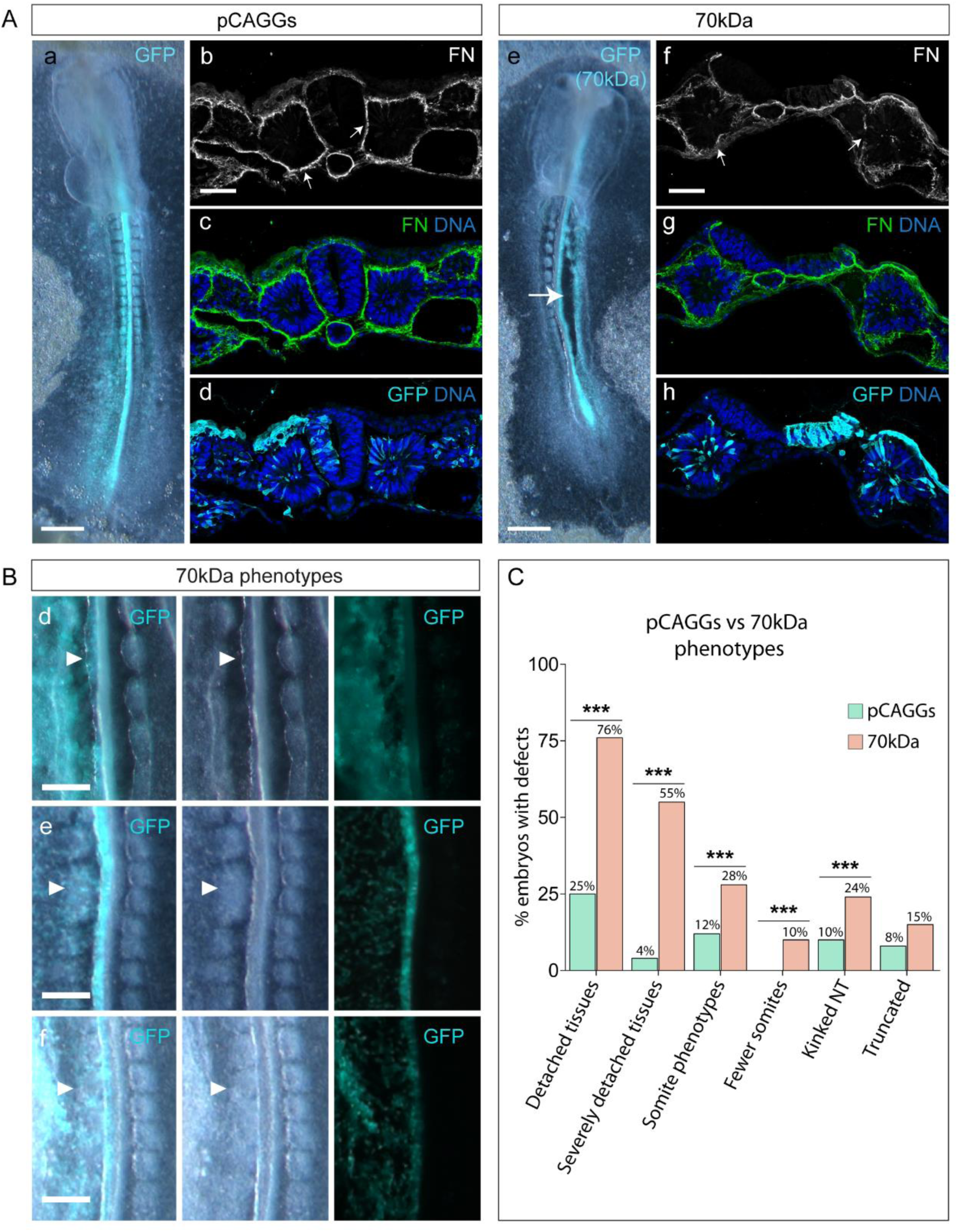
Embryos electroporated with the 70kDa construct exhibit numerous morphological defects. (**A**) Representative images of the morphology of embryos electroporated with either pCAGGs only (a) or 70kDa, the latter being a severe phenotype (e). Severe phenotypes included severely detached tissues (e, arrow) and a truncated A-P axis. Transverse sections of pCAGGs- (b-d) and 70kDa-electroporared embryos (f-h) immunostained for fibronectin (b-c, f-g), GFP (d, h) and stained for DNA (c-d, g-h). Electroporated side is on left (a-d) or right (e-h). Detachment of tissues is clearly visible in 70kDa-electroporated embryos (e), which is accompanied by a severe disruption in the fibronectin matrix (compare b-c with f-g, arrows). **(B)** Close up of embryos electroporated with 70kDa showing kinked neural tube (d, arrowheads), fused somites (e, arrowheads) and fewer somites on the electroporated side (f, arrowheads). Electroporated sides are on left. Ventral view and rostral on top. **(C)** Percentage of pCAGGS- (green bars) and 70kDa-electroporated (pink bars) embryos with morphological defects, including detached (pCAGGs: 39/154, 70kDa: 97/144) and severely detached tissues (pCAGGs: 6/154, 70kDa: 79/144), kinked neural tube (pCAGGs: 15/154, 70kDa: 34/144), truncated A-P axis (pCAGGs: 13/154, 70kDa: 21/144), abnormal somite morphology (pCAGGs: 18/154, 70kDa: 41/144), and fewer somites on the electroporated side compared to the control non-electroporated side (pCAGGs: 0/154, 70kDa: 15/144). p values were calculated using a Chi-square test. ***p<0.001. FN – fibronectin. Scale bars: (A, a, e; C, a-c) 500 µm, (C, d-f) 200 µm, (A, b-d, f-h) 50 µm.

**Supplementary Figure 7.**
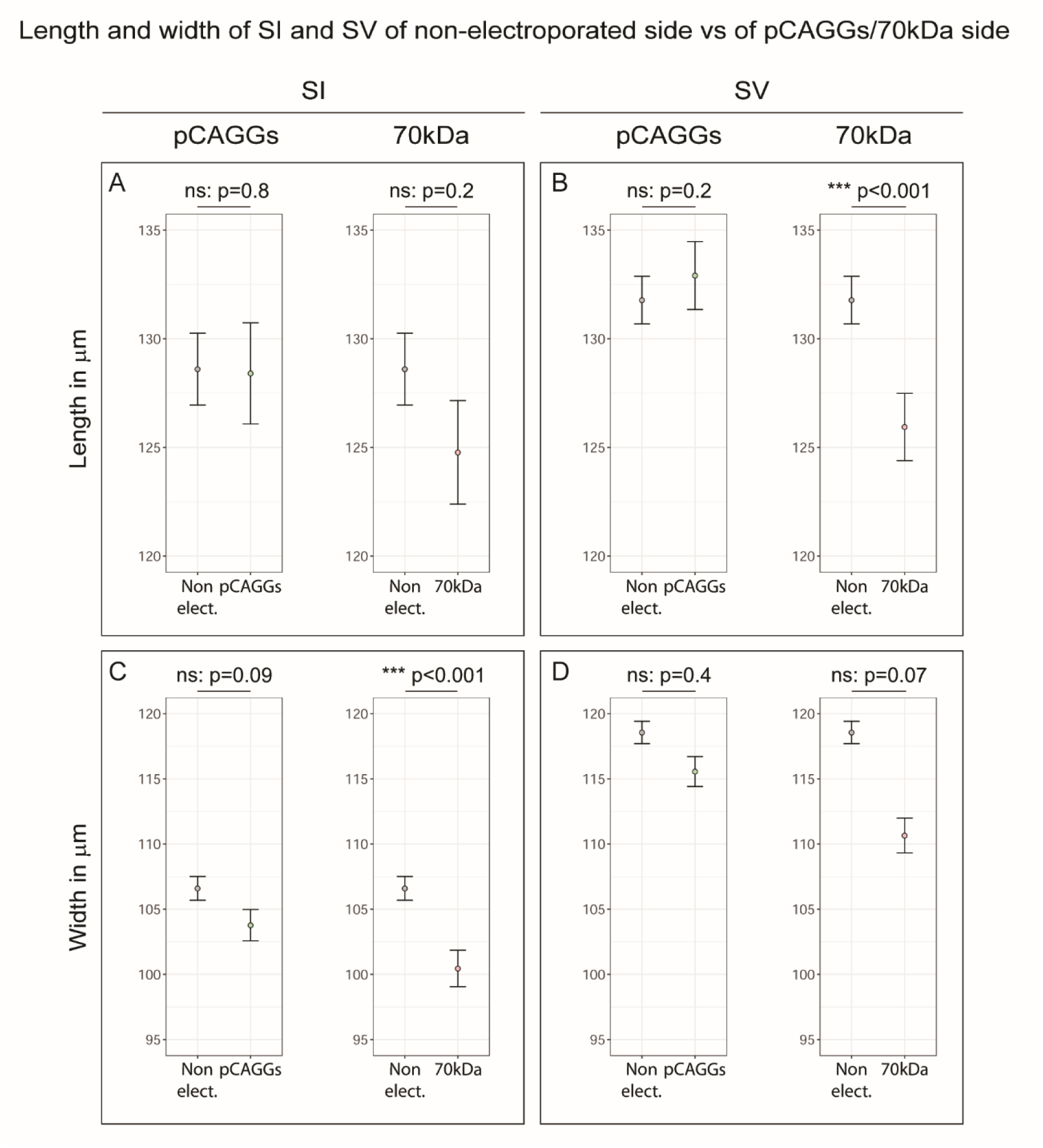
Quantification of somite length and width in pCAGGs- and 70kDa-electroporated embryos. (**A-D**) Length of SI (A) and SV (B) and width of SI (C) and SV (D) from pCAGGS- and 70kDa-electroporated embryos compared to the control non-electroporated side. The measurements were made on images from whole mount embryos. Somites from pCAGGs-electroporated embryos did not show a significant difference in either length or width between electroporated vs non-electroporated sides (n=151), but the width of SI and length of SV of the electroporated side of 70kDa-treated embryos were significantly smaller than that of the contralateral non-electroporated control side (n=143). Bars represent the standard error of the mean. p values were calculated using a nested ANOVA. ns – not significant; *** p<0.001.

## Notes

#### Summary of Updates

The text of the manuscript has been slightly revised.

## References

1. Aulehla A, Pourquié O. 2010. Signaling gradients during paraxial mesoderm development. Cold Spring Harb Perspect Biol 2:1–17. doi:10.1101/cshperspect.a000869

2. Bailey C, Dale K. 2015. Somitogenesis in Vertebrate Development.ELS. John Wiley & Sons, Ltd: Chichester. Chichester, UK: John Wiley & Sons, Ltd. pp. 1–15. doi:10.1002/9780470015902.a0003820.pub2

3. Barriga EH, Franze K, Charras G, Mayor R. 2018. Tissue stiffening coordinates morphogenesis by triggering collective cell migration in vivo. Nature 554:523–527. doi:10.1038/nature25742

4. Barrios A, Poole RJ, Durbin L, Brennan C, Holder N, Wilson SW. 2003. Eph/Ephrin signaling regulates the mesenchymal-to-epithelial transition of the paraxial mesoderm during somite morphogenesis. Curr Biol 13:1571–82. doi:10.1016/j.cub.2003.08.030

5. Bénazéraf B, Beaupeux M, Tchernookov M, Wallingford A, Salisbury T, Shirtz A, Shirtz A, Huss D, Pourquié O, François P, Lansford R. 2017. Multi-scale quantification of tissue behavior during amniote embryo axis elongation. Development 144:4462–4472. doi:10.1242/dev.150557

6. Bénazéraf B, Francois P, Baker RE, Denans N, Little CD, Pourquié O. 2010. A random cell motility gradient downstream of FGF controls elongation of an amniote embryo. Nature 466:248– 252. doi:10.1038/nature09151

7. Benito-Jardón M, Klapproth S, Gimeno-LLuch I, Petzold T, Bharadwaj M, Müller DJ, Zuchtriegel G, Reichel CA, Costell M. 2017. The fibronectin synergy site re-enforces cell adhesion and mediates a crosstalk between integrin classes. Elife 6:1–24. doi:10.7554/elife.22264

8. Brunet T, Bouclet A, Ahmadi P, Mitrossilis D, Driquez B, Brunet A-C, Henry L, Serman F, Béalle G, Ménager C, Dumas-Bouchiat F, Givord D, Yanicostas C, Le-Roy D, Dempsey NM, Plessis A, Farge E. 2013. Evolutionary conservation of early mesoderm specification by mechanotransduction in Bilateria. Nat Commun 4:2821. doi:10.1038/ncomms3821

9. Buchberger A, Seidl K, Klein C, Eberhardt H, Arnold HH. 1998. cMeso-1, a novel bHLH transcription factor, is involved in somite formation in chicken embryos. Dev Biol 199:201– 15. doi:10.1006/dbio.1998.8919

10. Burute M, Thery M. 2012. Spatial segregation between cell–cell and cell–matrix adhesions. Curr Opin Cell Biol 24:628–636. doi:10.1016/j.ceb.2012.07.003

11. Campbell ID, Humphries MJ. 2011. Integrin structure, activation, and interactions. Cold Spring Harb Perspect Biol 3:a004994–a004994. doi:10.1101/cshperspect.a004994

12. Chan CJ, Heisenberg C-P, Hiiragi T. 2017. Coordination of morphogenesis and cell-fate specification in development. Curr Biol 27:R1024–R1035. doi:10.1016/j.cub.2017.07.010

13. Chapman SC, Collignon J, Schoenwolf GC, Lumsden A. 2001. Improved method for chick whole-embryo culture using a filter paper carrier. Dev Dyn 220:284–9. doi:10.1002/1097- 0177(20010301)220:3<284::AID-DVDY1102>3.0.CO;2-5

14. Charras G, Yap AS. 2018. Tensile Forces and Mechanotransduction at Cell–Cell Junctions. Curr Biol 28:R445–R457. doi:10.1016/j.cub.2018.02.003

15. Chernoff EA, Hilfer SR. 1982. Calcium dependence and contraction in somite formation. Tissue Cell 14:435–49. doi:10.1016/0040-8166(82)90038-6

16. Christ B, Huang R, Scaal M. 2007. Amniote somite derivatives. Dev Dyn 236:2382–2396. doi:10.1002/dvdy.21189

17. Danen EHJ, Sonneveld P, Brakebusch C, Fässler R, Sonnenberg A. 2002. The fibronectin-binding integrins α5β1 and αvβ3 differentially modulate RhoA-GTP loading, organization of cell matrix adhesions, and fibronectin fibrillogenesis. J Cell Biol 159:1071–1086. doi:10.1083/jcb.200205014

18. Dequéant M-L, Glynn E, Gaudenz K, Wahl M, Chen J, Mushegian A, Pourquié O. 2006. A complex oscillating network of signaling genes underlies the mouse segmentation clock. Science 314:1595–8. doi:10.1126/science.1133141

19. Drake CJ, Davis LA, Hungerford JE, Little CD. 1992. Perturbation of beta 1 integrin-mediated adhesions results in altered somite cell shape and behavior. Dev Biol 149:327–38. doi:10.1016/0012-1606(92)90288-R

20. Drake CJ, Little CD. 1991. Integrins play an essential role in somite adhesion to the embryonic axis. Dev Biol 143:418–21. doi:10.1016/0012-1606(91)90092-H

21. Friedland JC, Lee MH, Boettiger D. 2009. Mechanically activated integrin switch controls alpha5beta1 function. Science 323:642–4. doi:10.1126/science.1168441

22. George EL, Georges-Labouesse EN, Patel-King RS, Rayburn H, Hynes RO. 1993. Defects in mesoderm, neural tube and vascular development in mouse embryos lacking fibronectin. Development 119:1079–91.

23. Georges-Labouesse EN, George EL, Rayburn H, Hynes RO. 1996. Mesodermal development in mouse embryos mutant for fibronectin. Dev Dyn 207:145–56. doi:10.1002/(SICI)1097-0177(199610)207:2<145::AID-AJA3>3.0.CO;2-H

24. Girós A, Grgur K, Gossler A, Costell M. 2011. α5β1 integrin-mediated adhesion to fibronectin is required for axis elongation and somitogenesis in mice. PLoS One 6:e22002. doi:10.1371/journal.pone.0022002

25. Goh KL, Yang JT, Hynes RO. 1997. Mesodermal defects and cranial neural crest apoptosis in alpha5 integrin-null embryos. Development 124:4309–19.

26. Gomes de Almeida P, Pinheiro GG, Nunes AM, Gonçalves AB, Thorsteinsdóttir S. 2016. Fibronectin assembly during early embryo development: A versatile communication system between cells and tissues. Dev Dyn 245:520–35. doi:10.1002/dvdy.24391

27. Gordon WR, Zimmerman B, He L, Miles LJ, Huang J, Tiyanont K, McArthur DG, Aster JC, Perrimon N, Loparo JJ, Blacklow SC. 2015. Mechanical Allostery: Evidence for a Force Requirement in the Proteolytic Activation of Notch. Dev Cell 33:729–36. doi:10.1016/j.devcel.2015.05.004

28. Hamburger V, Hamilton HL. 1992. A series of normal stages in the development of the chick embryo. Dev Dyn 195:231–272. doi:10.1002/aja.1001950404

29. Henrique D, Adam J, Myat A, Chitnis A, Lewis J, Ish-Horowicz D. 1995. Expression of a Delta homologue in prospective neurons in the chick. Nature 375:787–790. doi:10.1038/375787a0

30. Hiramatsu R, Matsuoka T, Kimura-Yoshida C, Han S-W, Mochida K, Adachi T, Takayama S, Matsuo I. 2013. External Mechanical Cues Trigger the Establishment of the Anterior-Posterior Axis in Early Mouse Embryos. Dev Cell 27:131–144. doi:10.1016/j.devcel.2013.09.026

31. Horton ER, Humphries JD, James J, Jones MC, Askari JA, Humphries MJ. 2016. The integrin adhesome network at a glance. J Cell Sci 129:4159–4163. doi:10.1242/jcs.192054

32. Hubaud A, Pourquié O. 2014. Signalling dynamics in vertebrate segmentation. Nat Rev Mol Cell Biol 15:709–721. doi:10.1038/nrm3891

33. Hubaud A, Regev I, Mahadevan L, Pourquié O. 2017. Excitable dynamics and Yap-dependent mechanical cues drive the segmentation clock. Cell 171:668–682.e11. doi:10.1016/j.cell.2017.08.043

34. Hunter GL, He L, Perrimon N, Charras G, Giniger E, Baum B. 2019. A role for actomyosin contractility in Notch signaling. BMC Biol 17:12. doi:10.1186/s12915-019-0625-9

35. Huveneers S, Truong H, Fässler R, Sonnenberg A, Danen EHJ. 2008. Binding of soluble fibronectin to integrin alpha5 beta1 - link to focal adhesion redistribution and contractile shape. J Cell Sci 121:2452–62. doi:10.1242/jcs.033001

36. Jouve C, Palmeirim I, Henrique D, Beckers J, Gossler A, Ish-Horowicz D, Pourquié O. 2000. Notch signalling is required for cyclic expression of the hairy-like gene HES1 in the presomitic mesoderm. Development 127:1421–1429. doi:10.1016/S0092-8674(00)80451-1

37. Jülich D, Cobb G, Melo AM, McMillen P, Lawton AK, Mochrie SGJ, Rhoades E, Holley SA. 2015. Cross-scale integrin regulation organizes ECM and tissue topology. Dev Cell 34:33–44. doi:10.1016/j.devcel.2015.05.005

38. Jülich D, Geisler R, Holley SA, Tübingen 2000 Screen Consortium. 2005. Integrinalpha5 and delta/notch signaling have complementary spatiotemporal requirements during zebrafish somitogenesis. Dev Cell 8:575–86. doi:10.1016/j.devcel.2005.01.016

39. Kocsis E, Trus BL, Steer CJ, Bisher ME, Steven AC. 1991. Image averaging of flexible fibrous macromolecules: the clathrin triskelion has an elastic proximal segment. J Struct Biol 107:6–14. doi:10.1093/bioinformatics/btp184

40. Koshida S, Kishimoto Y, Ustumi H, Shimizu T, Furutani-Seiki M, Kondoh H, Takada S. 2005. Integrin alpha5-dependent fibronectin accumulation for maintenance of somite boundaries in zebrafish embryos. Dev Cell 8:587–98. doi:10.1016/j.devcel.2005.03.006

41. Kragtorp KA, Miller JR. 2007. Integrin alpha5 is required for somite rotation and boundary formation in Xenopus. Dev Dyn 236:2713–20. doi:10.1002/dvdy.21280

42. Lauschke VM, Tsiairis CD, François P, Aulehla A. 2013. Scaling of embryonic patterning based on phase-gradient encoding. Nature 493:101–105. doi:10.1038/nature11804

43. Lawton AK, Nandi A, Stulberg MJ, Dray N, Sneddon MW, Pontius W, Emonet T, Holley SA. 2013. Regulated tissue fluidity steers zebrafish body elongation. Development 140:573–582. doi:10.1242/dev.090381

44. Luca VC, Kim BC, Ge C, Kakuda S, Wu D, Roein-Peikar M, Haltiwanger RS, Zhu C, Ha T, Garcia KC. 2017. Notch-Jagged complex structure implicates a catch bond in tuning ligand sensitivity. Science (80-) 355:1320–1324. doi:10.1126/science.aaf9739

45. Mao Y, Schwarzbauer JE. 2005. Fibronectin fibrillogenesis, a cell-mediated matrix assembly process. Matrix Biol 24:389–399. doi:10.1016/j.matbio.2005.06.008

46. Marek M, Kubícek M. 1981. Morphogen pattern formation and development in growth. Bull Math Biol 43:259–70.

47. Marrese M, Antonovaite N, Nelemans BKA, Smit TH, Iannuzzi D. 2019. Micro-indentation and optical coherence tomography for the mechanical characterization of embryos: Experimental setup and measurements on chicken embryos. Acta Biomater. doi:10.1016/j.actbio.2019.07.056

48. Martins GG, Rifes P, Amândio R, Rodrigues G, Palmeirim I, Thorsteinsdóttir S. 2009. Dynamic 3D cell rearrangements guided by a fibronectin matrix underlie somitogenesis. PLoS One 4:e7429. doi:10.1371/journal.pone.0007429

49. Masamizu Y, Ohtsuka T, Takashima Y, Nagahara H, Takenaka Y, Yoshikawa K, Okamura H, Kageyama R. 2006. Real-time imaging of the somite segmentation clock: Revelation of unstable oscillators in the individual presomitic mesoderm cells. Proc Natl Acad Sci 103:1313–1318. doi:10.1073/pnas.0508658103

50. McKeown-Longo PJ, Mosher DF. 1985. Interaction of the 70,000-mol-wt amino-terminal fragment of fibronectin with the matrix-assembly receptor of fibroblasts. J Cell Biol 100:364–74. doi:10.1083/jcb.100.2.364

51. Meloty-Kapella L, Shergill B, Kuon J, Botvinick E, Weinmaster G. 2012. Notch Ligand Endocytosis Generates Mechanical Pulling Force Dependent on Dynamin, Epsins, and Actin. Dev Cell 22:1299–1312. doi:10.1016/j.devcel.2012.04.005

52. Merle T, Farge E. 2018. Trans-scale mechanotransductive cascade of biochemical and biomechanical patterning in embryonic development: the light side of the force. Curr Opin Cell Biol 55:111–118. doi:10.1016/j.ceb.2018.07.003

53. Mongera A, Rowghanian P, Gustafson HJ, Shelton E, Kealhofer DA, Carn EK, Serwane F, Lucio AA, Giammona J, Campàs O. 2018. A fluid-to-solid jamming transition underlies vertebrate body axis elongation. Nature 561:401–405. doi:10.1038/s41586-018-0479-2

54. Morimoto M, Takahashi Y, Endo M, Saga Y. 2005. The Mesp2 transcription factor establishes segmental borders by suppressing Notch activity. Nature 435:354–359. doi:10.1038/nature03591

55. Morin-Kensicki EM, Boone BN, Howell M, Stonebraker JR, Teed J, Alb JG, Magnuson TR, O’Neal W, Milgram SL. 2006. Defects in Yolk Sac Vasculogenesis, Chorioallantoic Fusion, and Embryonic Axis Elongation in Mice with Targeted Disruption of Yap65. Mol Cell Biol 26:77– 87. doi:10.1128/MCB.26.1.77-87.2006

56. Mui KL, Chen CS, Assoian RK. 2016. The mechanical regulation of integrin–cadherin crosstalk organizes cells, signaling and forces. J Cell Sci 129:1093–1100. doi:10.1242/jcs.183699

57. Nakajima Y, Morimoto M, Takahashi Y, Koseki H, Saga Y. 2006. Identification of Epha4 enhancer required for segmental expression and the regulation by Mesp2. Development 133:2517– 25. doi:10.1242/dev.02422

58. Newell-Litwa KA, Horwitz R, Lamers ML. 2015. Non-muscle myosin II in disease: mechanisms and therapeutic opportunities. Dis Model Mech 8:1495–515. doi:10.1242/dmm.022103

59. Niwa Y, Shimojo H, Isomura A, Gonzalez A, Miyachi H, Kageyama R. 2011. Different types of oscillations in Notch and Fgf signaling regulate the spatiotemporal periodicity of somitogenesis. Genes Dev 25:1115–1120. doi:10.1101/gad.2035311

60. Palmeirim I, Dubrulle J, Henrique D, Ish-Horowicz D, Pourquié O. 1998. Uncoupling segmentation and somitogenesis in the chick presomitic mesoderm. Dev Genet 23:77–85. doi:10.1002/(SICI)1520-6408(1998)23:1<77::AID-DVG8>3.0.CO;2-3

61. Palmeirim I, Henrique D, Ish-Horowicz D, Pourquié O. 1997. Avian hairy gene expression identifies a molecular clock linked to vertebrate segmentation and somitogenesis. Cell 91:639–648. doi:10.1016/S0092-8674(00)80451-1

62. Piccolo S, Dupont S, Cordenonsi M. 2014. The Biology of YAP/TAZ: Hippo Signaling and Beyond. Physiol Rev 94:1287–1312. doi:10.1152/physrev.00005.2014

63. Pierschbacher MD, Ruoslahti E. 1984. Cell attachment activity of fibronectin can be duplicated by small synthetic fragments of the molecule. Nature 309:30–3.

64. Pourquié O, Tam PPL. 2001. A nomenclature for prospective somites and phases of cyclic gene expression in the presomitic mesoderm. Dev Cell 1:619–20. doi:10.1016/S1534- 5807(01)00082-X

65. Preibisch S, Saalfeld S, Tomancak P. 2009. Globally optimal stitching of tiled 3D microscopic image acquisitions. Bioinformatics 25:1463–1465. doi:10.1093/bioinformatics/btp184

66. Prieto D, Aparicio G, Machado M, Zolessi FR. 2015. Application of the DNA-specific stain methyl green in the fluorescent labeling of embryos. J Vis Exp e52769. doi:10.3791/52769

67. Rallis C, Pinchin SM, Ish-Horowicz D. 2010. Cell-autonomous integrin control of Wnt and Notch signalling during somitogenesis. Development 137:3591–3601. doi:10.1242/dev.050070

68. Rifes P, Carvalho L, Lopes C, Andrade RP, Rodrigues G, Palmeirim I, Thorsteinsdóttir S. 2007. Redefining the role of ectoderm in somitogenesis: a player in the formation of the fibronectin matrix of presomitic mesoderm. Development 134:3155–3165. doi:10.1242/dev.003665

69. Rifes P, Thorsteinsdóttir S. 2012. Extracellular matrix assembly and 3D organization during paraxial mesoderm development in the chick embryo. Dev Biol 368:370–381. doi:10.1016/j.ydbio.2012.06.003

70. Ringer P, Colo G, Fässler R, Grashoff C. 2017. Sensing the mechano-chemical properties of the extracellular matrix. Matrix Biol 64:6–16. doi:10.1016/j.matbio.2017.03.004

71. Saga Y. 2012. The mechanism of somite formation in mice. Curr Opin Genet Dev 22:331–338. doi:10.1016/j.gde.2012.05.004

72. Saga Y, Takeda H. 2001. The making of the somite: molecular events in vertebrate segmentation. Nat Rev Genet 2:835–845. doi:10.1038/35098552

73. Sato Y, Nagatoshi K, Hamano A, Imamura Y, Huss D, Uchida S, Lansford R. 2017. Basal filopodia and vascular mechanical stress organize fibronectin into pillars bridging the mesoderm-endoderm gap. Development 144:281–291. doi:10.1242/dev.141259

74. Sato Y, Sato Y, Kasai T, Kasai T, Nakagawa S, Nakagawa S, Tanabe K, Tanabe K, Watanabe T, Watanabe T, Kawakami K, Kawakami K, Takahashi Y, Takahashi Y. 2007. Stable integration and conditional expression of electroporated transgenes in chicken embryos. Dev Biol 305:616–24. doi:10.1016/j.ydbio.2007.01.043

75. Sato Y, Yasuda K, Takahashi Y. 2002. Morphological boundary forms by a novel inductive event mediated by Lunatic fringe and Notch during somitic segmentation. Development 129:3633–44.

76. Schiller HB, Hermann M-R, Polleux J, Vignaud T, Zanivan S, Friedel CC, Sun Z, Raducanu A, Gottschalk K-E, Théry M, Mann M, Fässler R. 2013. β1- and αv-class integrins cooperate to regulate myosin II during rigidity sensing of fibronectin-based microenvironments. Nat Cell Biol 15:625–636. doi:10.1038/ncb2747

77. Schwartz MA, DeSimone DW. 2008. Cell adhesion receptors in mechanotransduction. Curr Opin Cell Biol 20:551–556. doi:10.1016/j.ceb.2008.05.005

78. Shih NP, Francois P, Delaune EA, Amacher SL. 2015. Dynamics of the slowing segmentation clock reveal alternating two-segment periodicity. Development 142:1785–1793. doi:10.1242/dev.119057

79. Singh P, Carraher C, Schwarzbauer JE. 2010. Assembly of Fibronectin Extracellular Matrix. Annu Rev Cell Dev Biol 26:397–419. doi:10.1146/annurev-cellbio-100109-104020

80. Slack JMW. 1987. We have a morphogen! Nature 327:553–554. doi:10.1038/327553a0

81. Smutny M, Ákos Z, Grigolon S, Shamipour S, Ruprecht V, Čapek D, Behrndt M, Papusheva E, Tada M, Hof B, Vicsek T, Salbreux G, Heisenberg CP. 2017. Friction forces position the neural anlage. Nat Cell Biol 19:306–317. doi:10.1038/ncb3492

82. Straight AF, Cheung A, Limouze J, Chen I, Westwood NJ, Sellers JR, Mitchison TJ. 2003. Dissecting temporal and spatial control of cytokinesis with a myosin II Inhibitor. Science 299:1743–7. doi:10.1126/science.1081412

83. Takahashi Y, Inoue T, Gossler A, Saga Y. 2003. Feedback loops comprising Dll1, Dll3 and Mesp2, and differential involvement of Psen1 are essential for rostrocaudal patterning of somites. Development 130:4259–68. doi:10.1242/dev.00629

84. Takahashi Y, Koizumi K, Takagi A, Kitajima S, Inoue T, Koseki H, Saga Y. 2000. Mesp2 initiates somite segmentation through the Notch signalling pathway. Nat Genet 25:390–6. doi:10.1038/78062

85. Takeichi M. 2014. Dynamic contacts: rearranging adherens junctions to drive epithelial remodelling. Nat Rev Mol Cell Biol 15:397–410. doi:10.1038/nrm3802

86. Tiedemann H. 1976. Pattern formation in early developmental stages of amphibian embryos. J Embryol Exp Morphol 35:437–44.

87. Watanabe T, Sato Y, Saito D, Tadokoro R, Takahashi Y. 2009. EphrinB2 coordinates the formation of a morphological boundary and cell epithelialization during somite segmentation. Proc Natl Acad Sci 106:7467–7472. doi:10.1073/pnas.0902859106

88. Wolfenson H, Lavelin I, Geiger B. 2013. Dynamic regulation of the structure and functions of integrin adhesions. Dev Cell 24:447–58. doi:10.1016/j.devcel.2013.02.012

89. Wolfenson H, Yang B, Sheetz MP. 2019. Steps in Mechanotransduction Pathways that Control Cell Morphology. Annu Rev Physiol 81:585–605. doi:10.1146/annurev-physiol-021317-121245

90. Yang JT, Bader BL, Kreidberg Ja, Ullman-Culleré M, Trevithick JE, Hynes RO. 1999. Overlapping and independent functions of fibronectin receptor integrins in early mesodermal development. Dev Biol 215:264–277. doi:10.1006/dbio.1999.9451

91. Yang JT, Rayburn H, Hynes RO. 1993. Embryonic mesodermal defects in alpha 5 integrin-deficient mice. Development 119:1093–105.

92. Yarrow JC, Totsukawa G, Charras GT, Mitchison TJ. 2005. Screening for cell migration inhibitors via automated microscopy reveals a Rho-kinase inhibitor. Chem Biol 12:385–95. doi:10.1016/j.chembiol.2005.01.015

93. Zaidel-Bar R. 2013. Cadherin adhesome at a glance. J Cell Sci 126:373–378. doi:10.1242/jcs.111559

94. Zaidel-Bar R, Zhenhuan G, Luxenburg C. 2015. The contractome - a systems view of actomyosin contractility in non-muscle cells. J Cell Sci 128:2209–2217. doi:10.1242/jcs.170068

